# Functional Unknomics of the SAR11 clade using bioinformatics approaches

**DOI:** 10.64898/2025.12.11.693642

**Authors:** Satoshi Nishino, Kento Tominaga, Kimiho Omae, Teppei Deguchi, Koji Hamasaki, Susumu Yoshizawa, Yuki Nishimura

## Abstract

A substantial fraction of the genes in marine bacteria lack detectable sequence similarity to genes with known functions. These functionally uncharacterized genes—collectively referred to as the “unknome”—represent a largely unexplored genetic repertoire harboring insights into marine bacterial ecology. In this study, we explored the function of the unknome of SAR11 clade, the most abundant bacterial lineage in the ocean, with a particular focus on genes that provide insight into their ecology. Based on the COG classification, approximately 58% of SAR11 ortholog groups were classified as unknome. Among the SAR11 unknome, we successfully inferred the functions of 69 ortholog groups that are conserved in SAR11 clade by protein structure similarity searches and genomic context analyses. These ortholog groups include putative transporter components, supporting the current ecological understanding that SAR11 clade is specialized in substrate uptake to adapt to oligotrophic marine environments. Furthermore, structural analysis indicated that the DUF2237-containing protein, enriched in marine environments, has potential interactions with purine nucleotide-containing compounds. This may suggest the existence of unique nucleotide utilization mechanisms in marine bacteria. In addition, we found candidates of virus defense systems within the unknome, which demonstrates that diverse defense systems are present at least in one-third of the cultured SAR11 strains. The conservation of these viral defense systems, even within streamlined SAR11 genomes, suggests that they confer significant ecological advantages. Our exploration provided insights into the genetic basis of bottom-up processes (adaptation to oligotrophic environments) and top-down processes (antiviral defense strategy) contributing to ecological success of SAR11.

## Introduction

The SAR11 clade, also known as *Candidatus* Pelagibacterales, is the most abundant group of heterotrophic bacteria in marine environments, accounting for approximately 25% of all bacterioplankton cells in the ocean photic zone (1). Due to its high abundance, members of SAR11 clade play crucial roles in marine ecosystems and material cycles (2,3). The notable characteristics of SAR11 clade are their slow growth rate and highly streamlined genome, which typically lacks accessory genes such as antiviral defense systems and, occasionally, essential biosynthetic pathways (3–6). Due to their poor growth under laboratory conditions, culture-based studies of SAR11 clade are limited in achieving comprehensiveness and throughput, and the reasons for their ecological dominance are still not fully resolved.

The ecological success of SAR11 has been explored from both bottom-up and top-down perspectives. On the bottom-up side, their extremely resource-efficient lifestyle is thought to provide a strong advantage in oligotrophic marine environments. Several observations supports this view, including the genome streamlining (5,7), presence of multifunctional transporters and highly expressed solute binding-proteins (8–10), loss of motility and transcriptional regulation (5,7,11), and light-utilization by proteorhodopsin (12,13). In contrast, from a top-down perspective, SAR11 are considered to be less affected by predation and lysis, in line with the “Kill-the-Winner” hypothesis. According to the hypothesis, fast-growing species are expected to be more susceptible to viral infections, implying that the slow-growing SAR11 strains are less affected by viral infections (14,15). However, the isolation of Pelagiphages (16) and evidence of their frequent infections to SAR11 (17–20) indicate that virus-host interactions are more complex than initially assumed. The “King of the Mountain” (KoM) hypothesis offers one possible explanation, proposing that the high abundance of SAR11 leads to increased contact frequency, promoting frequent genetic recombination, which could influence viral susceptibility (21). The KoM hypothesis is supported by the high intraspecific recombination rate and the presence of highly diverse genomic regions that encode membrane proteins that could act as viral receptors, variation that may help SAR11 populations evade viral infection (22–24). However, there is still a lack of direct evidence for the presence of antiviral defense systems and their frequent horizontal gene transfer among SAR11 (3,20).

Currently, genome-based approaches play indispensable roles in microbiology, particularly for lineages that are difficult to cultivate, such as the SAR11 clade. However, a growing challenge in the field is the discovery of a large number of genes with no similarity to any functionally characterized sequences (25,26). The issue is particularly acute in marine metagenomes (25,27), suggesting that ecologically important genes may remain overlooked and annotated as hypothetical proteins. For example, the recent discoveries of diverse antiviral defense systems originating form previously uncharacterized genes (28) suggest that many important functions await discovery within this uncharacterized gene space. To overcome the limitation of sequence-based annotation, an increasing number of studies are adopting bioinformatics approaches that go beyond sequence similarity, including protein structure prediction and genomic context analysis (29–31). However, these cutting-edge approaches that could substantially enhance functional annotation have not been systematically applied to SAR11 clade.

Here, we focused on the functionally unknown genes, referred to as the ‘functional unknome’ (26), within SAR11 genomes and sought to predict their functions by combining bioinformatics approaches. We placed particular emphasis on genes that may be ecologically important for the SAR11 clade, including those related to substrate uptake, phototrophy, and virus-host interactions. By combining structure-based predictions, genomic context information, and environmental distribution patterns, we inferred potential functions for conserved components of the SAR11 unknome. Further, we systematically explored viral defense systems within the SAR11 unknome. Our findings demonstrate the importance of re-annotating genes with unknown functions to uncover hidden aspects of non-model marine lineages.

## Materials and Methods

### Collection and evaluation of the SAR11 genomes

We compiled the SAR11 genomes that were analyzed by Haro-Moreno et al. (2020) (32) and expanded the dataset with genomes from freshwater, brackish water, Eastern Pacific, and polar environments to better represent the under-sampled clades (**Table S1**) (33–36). The final dataset includes 321 genomes, mainly from cultured strains and single-amplified genomes (SAGs), as metagenome-assembled genomes (MAGs) are difficult to reconstruct for SAR11 due to high genomic diversity (Fig. S1) (37). Genome quality was assessed by using CheckM2 v1.0.2 (**Fig. S1A**) (38). Estimated genome size was calculated using Completeness and Contamination scores following the previous study (36), and GC content was calculated using Seqkit v2.4.0 (**Fig. S1B**) (39). Among the 253 genomes with >50% completeness and <10% contamination, we identified at least 184 species based on <95% ANI, as calculated using FastANI v1.34 (40).

### Phylogenomic classification of the SAR11 genomes

The concatenated alignment of 120 bacterial single-copy genes (5,036 sites;) was constructed using GTDB-Tk v2.1.0 classify-wf (41). Eight genomes were excluded because they had <10% of the alignment sites covered. The *Rickettsia bellii* genome (GCA_002078315.1) served as the outgroup following a previous study (32). The maximum-likelihood tree was inferred with IQ-TREE2 v2.2.0.3 using ModelFinder-selected Q.insect+F+I+R9 model, with 1,000 ultrafast bootstrap and 1,000 SH-aLRT replicates (42,43). The tree was visualized in R using ggplot2, ggtree, and ggtreeExtra (44–46). Subclades were assigned based on tree topology following the previous classifications (6,3,32,36); genomes falling outside known subclades remained unassigned. Although the phylogenetic positions of subclades IV and V are debated and are not classified within the *Ca.* Pelagibacterales in several previous studies (32,47–49), we treated them as part of SAR11 clade, considering their shared genomic and ecological characteristics (3).

### Functional annotation of ortholog groups and definition of the unknome

A total of 306,416 protein-coding genes in all 321 SAR11 genomes were predicted with prokka v1.14.6 (50), and clustered into ortholog groups (OGs) using SonicParanoid2 v2.0.3 (51). Transmembrane regions were predicted using TMHMM 2.0 (52), and proteins were classified as transmembrane (TM) if they met either of the following criteria: (i) ExpAA >10 aa and PredHel >1, or (ii) ExpAA >10 aa, PredHel = 1, and First60 <10. All coding sequences were annotated with COG-2020 and KEGG Orthology (KO) using COGclassifier v1.0.5 (https://github.com/moshi4/COGclassifier) and KofamScan v1.3.0 (53), respectively. The unknome genes were defined as those were not annotated by COGclassifier or assigned COGs belonging to category S (Function unknown), category R (General function prediction only); unknome OGs were those with >50% unknome members. The OGs distributions among each subclade were visualized with the UpSetR (54). Environmental distribution data were retrieved from Global Microbial Gene Catalog (GMGC) v1.0 for bacteriorhodopsin and core unknome OGs based on their COG IDs (55). The core unknome was defined as unknome OGs occurring in more than half of the SAR11 genomes. Only GMGC entries annotated with both habitat categories (e.g., marine, freshwater and human gut) and COG IDs were analyzed. Habitat proportions were calculated per OG, counting multiple habitat annotations as independent occurrences. To simplify classification, “human skin,” “human gut,” “human nose,” “human vagina”, and “human oral” were grouped as “human-associated”.

### Neighborhood OG analysis

Genomic contexts of all coding sequences were analyzed using GFF files generated by the prokka pipeline. Two genes were considered neighbors if they were located within five genes of each other on the same contig. Neighboring pairs that showed a reciprocal co-localization probability of ≥75% across at least ten genomes were retained and visualized as a network using igraph (56) and ggnetwork (57). Since our analysis focused only on a phylogenetically close lineage, we adopted a more conservative threshold than the previous study (29) to avoid false positives due to phylogenetic constraints. Note that pairs with highly unequal occurrence frequencies may not meet this criterion, even if the rarer OG always co-occurs with the more prevalent one. Neighboring pairs containing unknome OGs are manually checked using predicted structures by AlphaFold2, and the complex formation was evaluated by pDockQ and pDockQ2 scores (58–60).

### Collection and comparison of predicted protein structures

All protein structures of *Ca.* Pelagibacter ubique HTCC1062 strain and *Ca.* Pelagibacter sp. IMCC9063 strain were downloaded from the AlphaFold Protein Structure Database (61). To improve the structural prediction of the conserved 108 core unknome OGs, the structures were predicted with AlphaFold2 based on the core unknome proteins of HTCC1062 excluding the signal peptide regions predicted by DeepTMHMM ver 1.0.24 (62). Structural similarity search for the unknome proteins were performed using Foldseek version 7.04e0ec8 (63) against the Protein Data Bank (PDB), which was downloaded on September 4, 2023 (64).

### Meta-transcriptome data analysis of SAR11 genes

Sequence reads of the 509 meta-transcriptomes obtained in the Tara Ocean samples were mapped onto the 321 SAR11 genomes (**Table S2**) (65). We downloaded paired-end sequence data from NCBI SRA by SRA toolkit (https://github.com/ncbi/sra-tools), and then quality control was performed by using fastp with options -q 20 -n 10 -l 60 (66). The quality-controlled reads were mapped via bwa-mem2 (67). Transcripts per million (TPM) were calculated using a script from a previous study (https://github.com/yosuken/CountMappedReads2) (68).

### Analysis of DUF2237 domain-containing proteins

A total of 7,968 DUF2237 domain-containing proteins were retrieved from the Pfam database on April 21, 2023. These sequences were annotated with domain information using pfam_scan (69) and aligned using MAFFT v7.508 (70) in the auto mode. The alignment was visualized on the predicted monomer structure of the DUF2237-containing protein of *Ca.* P. ubique HTCC1062 strain using ConSurf-DB (71) and PyMOL v2.5.4 (72). The Predicted Aligned Error plot was visualized using a Python package (https://github.com/cddlab/alphafold3_tools).

The OceanDNA MAG catalog v1 (73) and the Pfam database were utilized to assess the phylogenetic distribution of DUF2237 genes. Genes annotated as COG3651 and COG5524 were identified as DUF2237-containing proteins and proteorhodopsin, respectively. A multiple sequence alignment of 120 marker genes from 7,696 representative MAGs was generated with the GTDB-Tk classify-wf. The resulting alignment contained 5,036 sites. Subsequently, a maximum likelihood phylogenetic tree was inferred with IQ-TREE2 (v2.2.0.3) using ModelFinder-selected Q.yeast+R10 model, with 1,000 SH-aLRT replicates. The phylogenetic distribution of DUF2237 and proteorhodopsin was visualized on the phylogenetic tree using ggplot2, ggtree, and ggtreeExtra (44–46).

The structure of the *Ca. P.* ubique HTCC1062 strain’s DUF2237 protein, predicted by AlphaFold2, was prepared in PDBQT format using MGL Tools v1.5.7 (74) for docking simulations. A ligand library was constructed by converting 258 compounds with biological roles from the KEGG Compound database (br8001) and seven nucleotide-containing molecules from the Protein Data Bank Japan (NAI, PO4, NDP, FAD, COA, C2E, NAD) into PDBQT format using Open Babel v3.1.0 (75) and MGL Tools **(Table S3)**. The grid box was defined as 25 × 20 × 20 Å to the pocket site of the DUF2237 protein predicted by PrankWeb 3 (76). All docking simulations were performed using Autodock vina v1.2.5 (77) with an exhaustiveness value of 100. The possibility of binding to ATP was evaluated using ATPbind (78).

### Defense system annotation

DefenseFinder (updated on June 5, 2024) was used to identify potential antiviral defense genes (79). To check for the completeness of defense systems, neighboring genes of each defense gene found by DefenseFinder were manually annotated and compared to known defense genes registered to DefenseFinder wiki (80) by genomic context, sequence alignment using MAFFT, and structure similarity search using AlphaFold2 and Foldseek.

## Result

### Collection and phylogenomic analysis of SAR11 genomes

We collected 321 SAR11 genomes representing five subclades (**Table S1**). Subclade classification based on a maximum-likelihood tree inferred from GTDB marker genes was largely consistent with those inferred from 16S rRNA gene and 232 conserved single-copy genes in previous studies (3,6,32) **(Fig. 1A)**. Subclade V exhibited larger genome size than other subclades **(Fig. S1B)**, consistent with its higher metabolic potential (49). The 306,416 protein-coding genes were clustered into 5,603 OGs using SonicParanoid2 (81). To estimate the unknome fraction, all genes were annotated using the KO and the COG databases. As a result, 173,549 and 257,412 genes were assigned to KO and COG, respectively **(Fig. 1B)**. At the OG level, 1,286 OGs (23%) and 2,683 OGs (48%) were assigned KO and COG annotation, respectively **(Fig. 1CD and Table S4)**.

**Figure 1.**
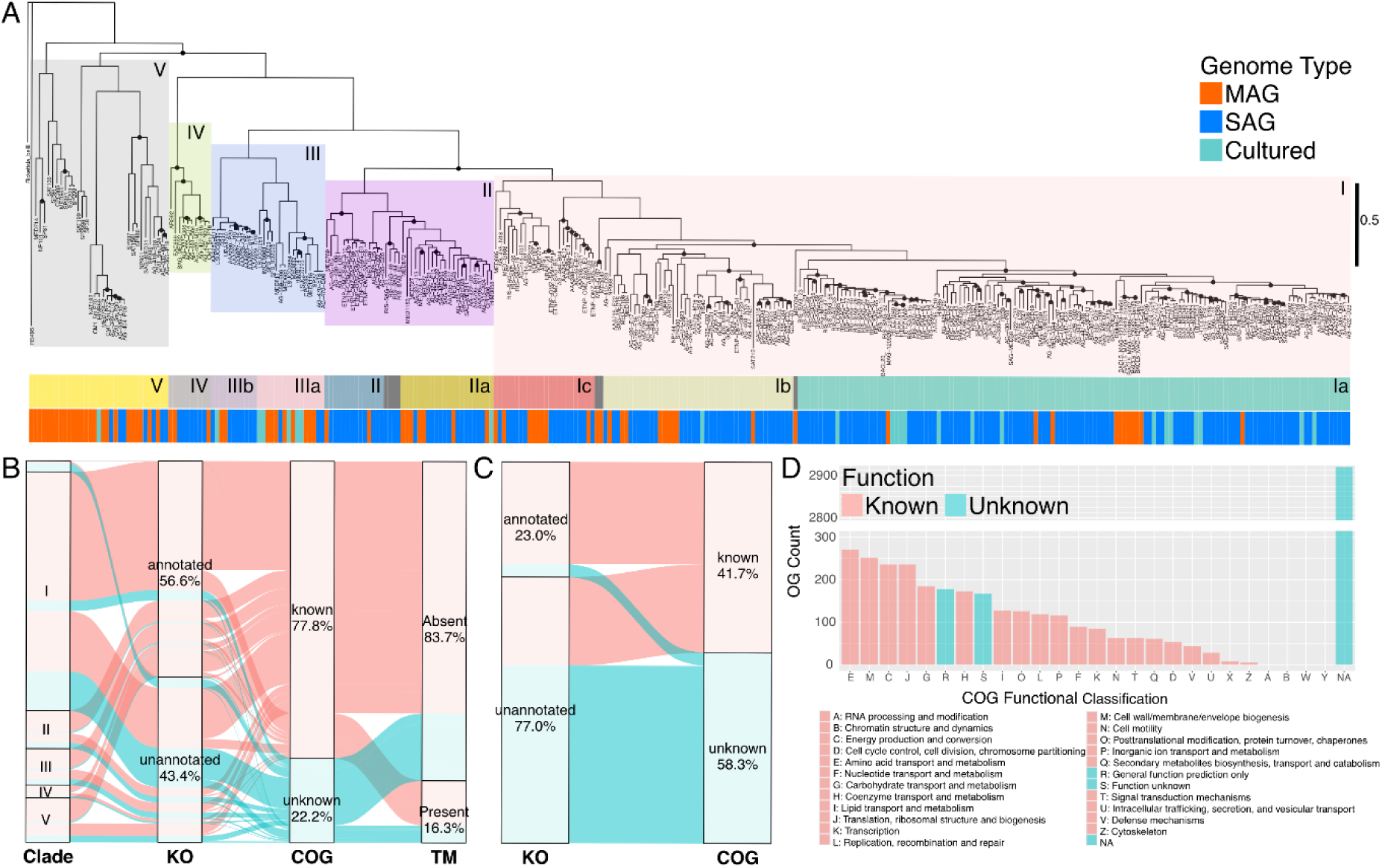
Characterization of the Unknome from a phylogenetically comprehensive SAR11 genome dataset. **(A)** Maximum likelihood phylogenetic tree of SAR11 genomes. The color coding in the phylogenetic tree indicates five subclade classifications. The colors below provides more detailed subclade classifications and types of the genomes, that is MAG (red), SAG (blue), or pure culture (light blue). The dots on the tree show UltraFast Bootstrap values > 95%. **(B,C)** Sankey plots summarize the annotation profiles of all genes/OGs and transmembrane region prediction results. Blue shows the unknome genes/OGs. TM indicates transmembrane proteins predicted using TMHMM 2.0 (see Methods for details). **(D)** COG annotation results for all OGs. NA denotes sequences not annotated in the COG database. Blue bars show unknome genes in this study.

We next classified these OGs according to their conservation within the five SAR11 subclades. The largest set combination corresponded to OGs conserved across all subclades (968 OGs), exceeding those unique to any single subclade **(Fig. S2A)**. This pattern is consistent with the streamlined SAR11 genomes, in which the core genome constitutes a large proportion (5,6). In contrast, most OGs with unknown functions were specific to clade I or clade V (790 and 557 OGs; **Fig. S2B)**, reflecting diversity of clade I and the large genome size of clade V (49). Additionaly, 192 unknown-function OGs specific to clade III, a brackish and freshwater clade, may contribute to the low-salinity adaptation **(Fig. S2B)**.

Among the COG-assigned genes, categories E (amino acid transport and metabolism, 271 OGs), M (cell wall and plasma membrane biosynthesis, 252 OGs), C (energy production, 236 OGs), and J (translation and ribosome-related, 236 OGs) and were the most frequently observed in the SAR11 clade **(Fig. 1D)**. The ordering of these categories, such as category M exceeding category K (transcription), contrasts with that of *E. coli* (82). This pattern reflects their reduced transcriptional regulation, a feature of genome streamlining (5–7). KO-assigned genes were fewer than COG-assigned genes **(Fig. 1BC)** and included functionally unknown genes, such as those containing domains of unknown function (DUF).

### Definition and characteristics of the ‘unknome’

Because some KOs lack functional annotation, we defined the SAR11 unknome based on COG classifications. It comprised category S, R, and OGs without COG assignment **(Fig. 1B)**, totaling 3,255 OGs (58.1%) **(Fig. 1D)**. The unknome genes are typically short and AT-rich **(Fig. S3AB)**. As in *E. coli* (83), these unknome genes show a slightly higher proportion of TM proteins than functionally known genes **(Fig. 1C, Fisher’s exact test p < 2.2e-16)**. Tara Oceans metatranscriptomes confirmed the unknome OGs are expressed in situ, with only slightly lower expression than functionally known genes **(Fig. S3C and Fig. S4, Wilcoxon rank sum test p < 2.2e-16)**.

### Functional prediction of the unknome based on neighboring gene functions and predicted protein structures

Next, we predict the functions of the unknome using two bioinformatics approaches. The first was neighborhood gene analysis, which predicts gene function from the genomic context because functionally related genes in prokaryotes often occur in close proximity in prokaryotic genomes (e.g., in operons or syntenic regions) (84,85). We defined neighboring OG pairs as those located within five genes of each other (details in Methods). We then constructed a neighboring OG network containing 474 OGs and 620 edges **(Fig. 2A)**. Functionally related OGs formed connected components (**Fig. 2A**), the largest of which corresponded to the ribosomal protein operon.

**Figure 2.**
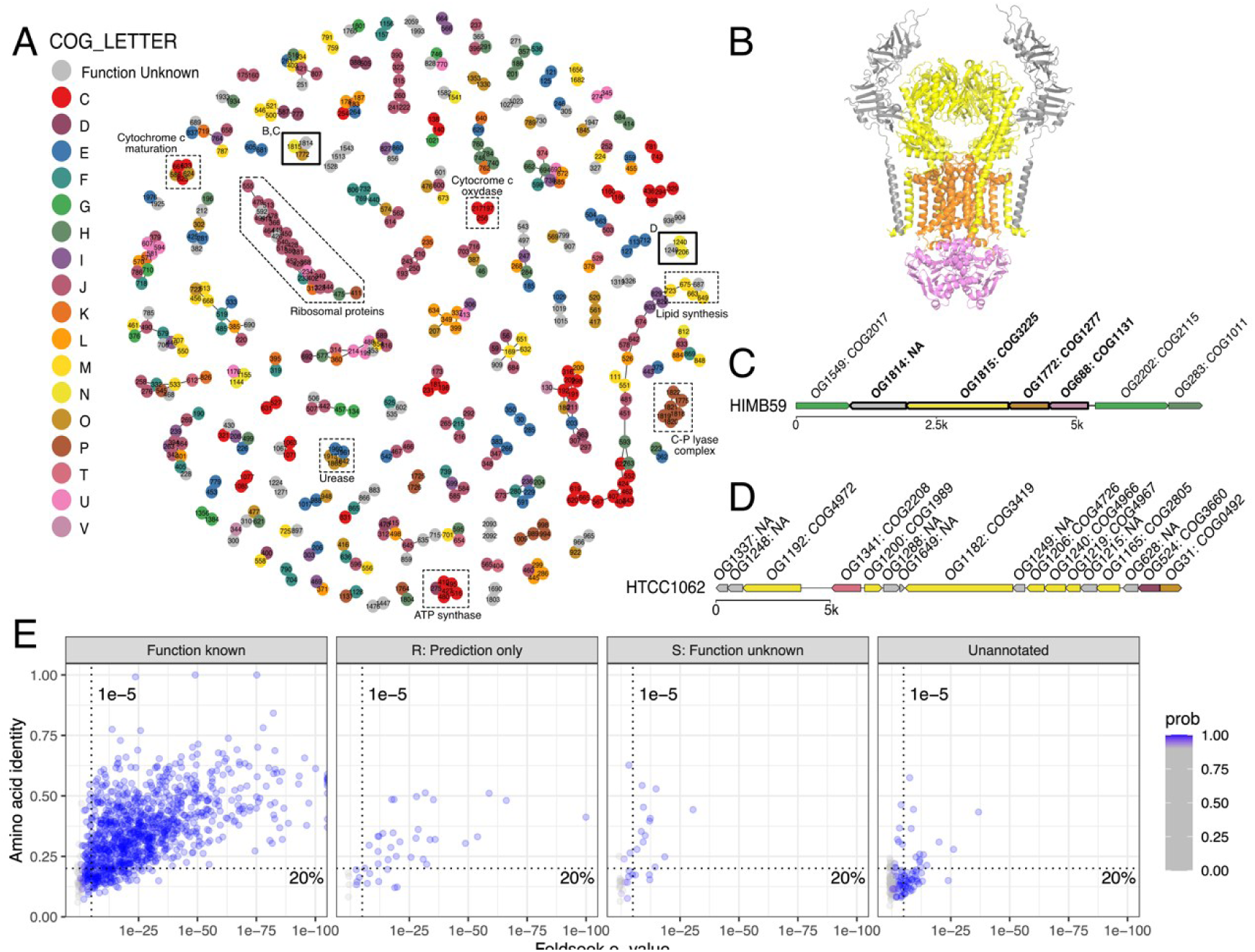
Function prediction based on genomic context and protein structural similarity search. **(A)** Summary of neighboring gene analysis. Network nodes represent each OGs, and colors indicate the COG functional categories. Pairs with a co-localization probability ≧75% within five genes upstream and downstream on the same contig are connected by edges. Several known operons are highlighted with dashed outlines as examples. **(B)** Predicted protein complex structure of OG688, OG1772, OG1815, and OG1549 using AlphaFold-multimer. The complex forms an octameric assembly composed of four homodimers. The colors show the COG functional category. **(C, D)** Genome context around each network component in panel A. The genes outlined in bold were used for protein structure prediction in panel B. **(E)** Scatter plots of all *Ca.* P. ubique HTCC1062 strain proteins compared with known protein structures in the PDB. The dot lines show e-value = 1e-5 and amino acid identity = 20%, highlighting potential remote homologs

Focusing on the connected components that include the unknome OGs (gray nodes), OG1814, a DUF4340-containing OG without a COG assignment, was conserved near the genes annotated as ABC transporter components, such as OG 1815, OG1772, and OG688. In HIMB59 (GCA_000299115.1), proteins from these four OGs were predicted to form a transporter complex with a pDcokQ2 score of 0.268, greater than the confidence threshold of 0.23 **(Fig. 2BC and Fig. S5AB, pDockQ2 > 0.268)**, suggesting that OG1814 function as a novel transporter subunit. Similarly, OG1249 (COG4696, a DUF2139-containing gene) was conserved within the type IV pilus operon, indicating a potential role as a minor pilus component (**Fig. 2D**). In total, 38 components containing both unknome and characterized OGs were identified. Although detailed functions remain unresolved for many, these patterns reveal functional links within the SAR11 clade—between unknome and known OGs, and among the unknome themselves.

We next applied protein structure prediction and structure similarity searches. Using *Ca.* P. ubique HTCC1062 (subclade Ia.3) and LD12 IMCC9063 (subclade IIIa.2) as the representatives, we compared predicted protein structures to those in the PDB. In HTCC1062, 1,212/1,328 proteins (91%) showed >90% probability of being homologous to a PDB entry, and 237 of these shared <20% sequence identity with their best hits (**Fig. 2E**). In IMCC9063 strain, 1,216/1,382 proteins (88%) showed high probability of structural homologs, 221 of them having <20% sequence identity **(Fig. S6**). Among unknome proteins, structurally similar proteins were found for 154/242 (67%) in HTCC1062 and 164/293 (56%) in IMCC9063. These results demonstrate that structural similarity searches effectively detect remote homologs within the SAR11 genomes.

### Function prediction of the core unknome OGs

Given that OGs widely conserved throughout the SAR11 clade likely play critical roles for their ecology, we further focused on the 108 core unknome OGs that are conserved across all five clades and present in >162 genomes (>50% of genomes in the dataset). By integrating information from neighboring genes, structurally similar proteins, and KO assignments, 87 of 108 OGs had at least one line of supporting information (**Fig. S7**). However, in many cases, the assigned KO showed no functional information, or the functions of the similar structures or neighboring genes were also unknome (**Supplemental Text**). Then, we performed an extensive literature search using gene and domain names registered in the COG, KO, and STRING databases to capture functional information not yet incorporated into current annotations. This search identified functional insights for 10 OGs, including several characterize in *Caulobacter crescentus*, a model alpha-proteobacterium (**Supplemental Text**). Overall, we found that 69 of 108 core unknome OGs have additional information or putative functions not registered in COG database **(Table S5 and Supplemental Text)**. These include predicted components of transporters, channels, or metabolic enzymes (e.g., OG228, OG305, OG365, OG448, OG497, OG814, and OG824), which may support the SAR11 clade’s adaptation to nutrient-limited environments (**Fig S8 and Supplemental Text**).

### Functional analysis of DUF2237, the unknown gene with a skewed distribution in the ocean

We next examined ecological distribution to infer their potential functions of core unknome OGs lacking functional information from any of the above analyses. We focused on the OGs enriched in marine environment, as such genes often underpin ecological success. A representative example is microbial rhodopsin (COG5524: bacteriorhodopsin) (**Fig. 3A**), which function as a photoreceptor, enabling the capture and utilization of light enagy in oligotrophic waters (86). The most marine-biased functionally unknown OG was COG3908, which contains the DUF2093 domain **(Fig. 3A)**. Although no further functional information was obtained, this OG was consistently located near the lipid synthesis operon in SAR11 clade (**Fig. 2A**).

**Figure 3.**
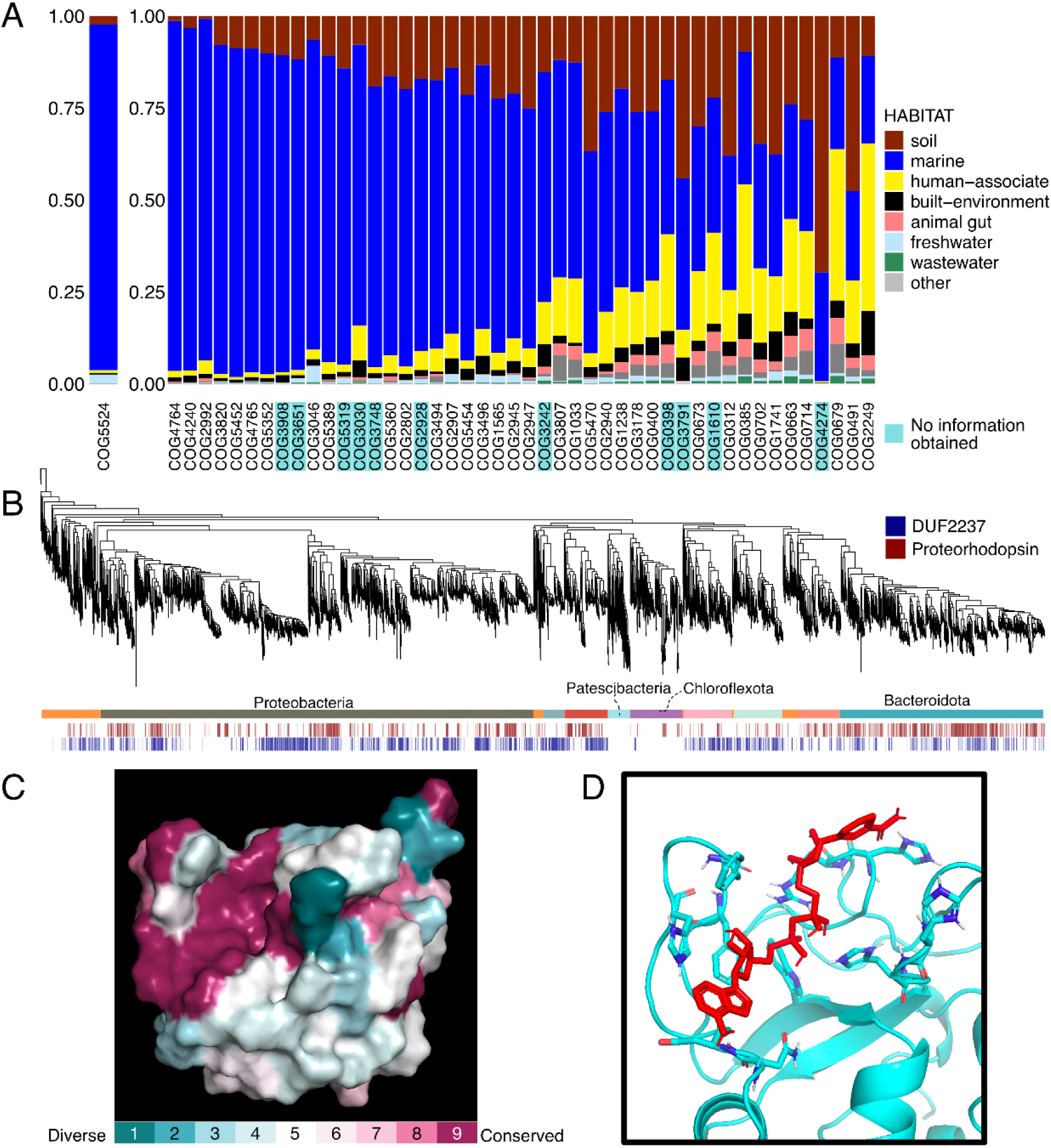
The core unknome distribution and functional analysis of DUF2237-containing protein. **(A)** Environmental distribution of bacteriorhodopsin (COG5524) and COG-assigned core unknomes based on GMGC. Bar plot colors indicate habitat, and the COG ID color indicate whether function could be inferred using the bioinformatics approaches. **(B)** Phylogenetic distribution of the DUF2237-containing gene and proteorhodopsin genes on representative bacterial genomes in the OceanDNA MAG catalog. The heatmap below indicates phylum, and the presence of DUF2237-containing gene and proteorhodopsin gene. **(C)** The amino acid conservation of 7,963 DUF2237-containing genes in the Pfam database, mapped onto the predicted structure of the DUF2237 protein from *Ca.* P. ubique HTCC1062. Conservation is color-coded from blue (<10%), to white (50%), to purple (>90%) based on the multiple sequence alignment. **(D)** Top docking pose of *P. ubique* HTCC1062 DUF2237 protein (light blue) and NADH (red).

The second most marine-biased OG was annotated as COG3651, which contains uncharacterized DUF2237 domain. DUF2237-containing genes have been reported to co-occur with the proteorhodopsin, a light-driven proton pumps, in marine *Flavobacteriia*, and a DUF2237 knockout in *Synechocystis* sp. PCC6803-P showed reduced phototaxis (87). DUF2237 has been suggested to have a function related to light reception or utilization. Although, three different structures of the DUF2237 protein had been experimentally solved (PDB 2LQ3, 3USH, and 4IOH), the mechanistic link to phototaxis remain unclear. Because light utilization is also ecologically important for SAR11 clade (13,88,11), we further analyzed this unknome protein.

Initially, we re-evaluated the association between proteorhodopsin and DUF2237-containing genes using OceanDNA MAG catalog (73) to validate the previous observation in marine *Flavobacteriia*. As in the previous study, DUF2237 genes significantly co-occurred with the proteorhodopsin **(Fisher’s exact test, p < 2.2e-16).** The high conservation of DUF2237 genes across most marine bacterial lineages, except *Patescibacteria* and *Chloroflexota* **(Fig. 3B)**, indicates their broad ecological importance.

The predicted structure of the DUF2237-containing protein (SAR11_1247) indicates dimeric assembly **(Fig. S9AB, pDockQ score = 0.314)**. Analysis of 7,968 DUF2237 domain-containing proteins in the Pfam database showed that this domain typically occurs as a single-domain protein **(Fig. S10AB)**. Mapping sequence conservation from these Pfam entries onto the structure revealed a highly conserved pocket region, likely a concave ligand-binding site **(Fig. 3C)**. This suggests a potential enzymatic role for DUF2237 in interacting with a specific ligand.

To identify potential ligands for the DUF2237 pocket, we performed docking simulations using KEGG “Compounds with biological roles”, excluding polysaccharides to avoid including macromolecules. Nucleotide-containing ligands, including NADH, c-di-GMP, and cAMP, showed the highest affinities for the DUF2237 protein from HTCC1062 strain (**Fig. 3D, Table S3**). ATP-binding prediction also supported nucleotide interaction (78) (Probability = 0.85). Because NADH and ATP participate in key metabolic pathways, and c-di-GMP and cAMP act as motility-related second messengers (89–92), these results suggest that the reduced phototaxis observed in the DUF2237 knockout mutant may arise not from a defective light response but rather from altered levels of small nucleotide-containing molecules that could mediate phototactic signaling.

### Antiviral defense genes on the SAR11 genomes

Given the numerous recent discoveries of defense genes within previously uncharacterized genes (28), we explored putative antiviral defense genes within the SAR11 genomes. Using DefenseFinder (80), we identified 43 defense systems, including restriction-modification (RM) and toxin-antitoxin systems **(Table S6)**. The RM systems were most common as previously reported (3,20). Excluding the well-characterized RM systems, 11 unknome genes showed putative defense roles **(Fig. S11)**. Furthermore, we examined genomic regions with only partial components of known systems and manually curated neighboring genes, as defense genes often cluster within specific loci. This led to the identification of 8 additional putative defense systems, bringing the total to 51 putative defense systems across 46 SAR11 genomes **(Fig. 4A)**.

**Figure 4.**
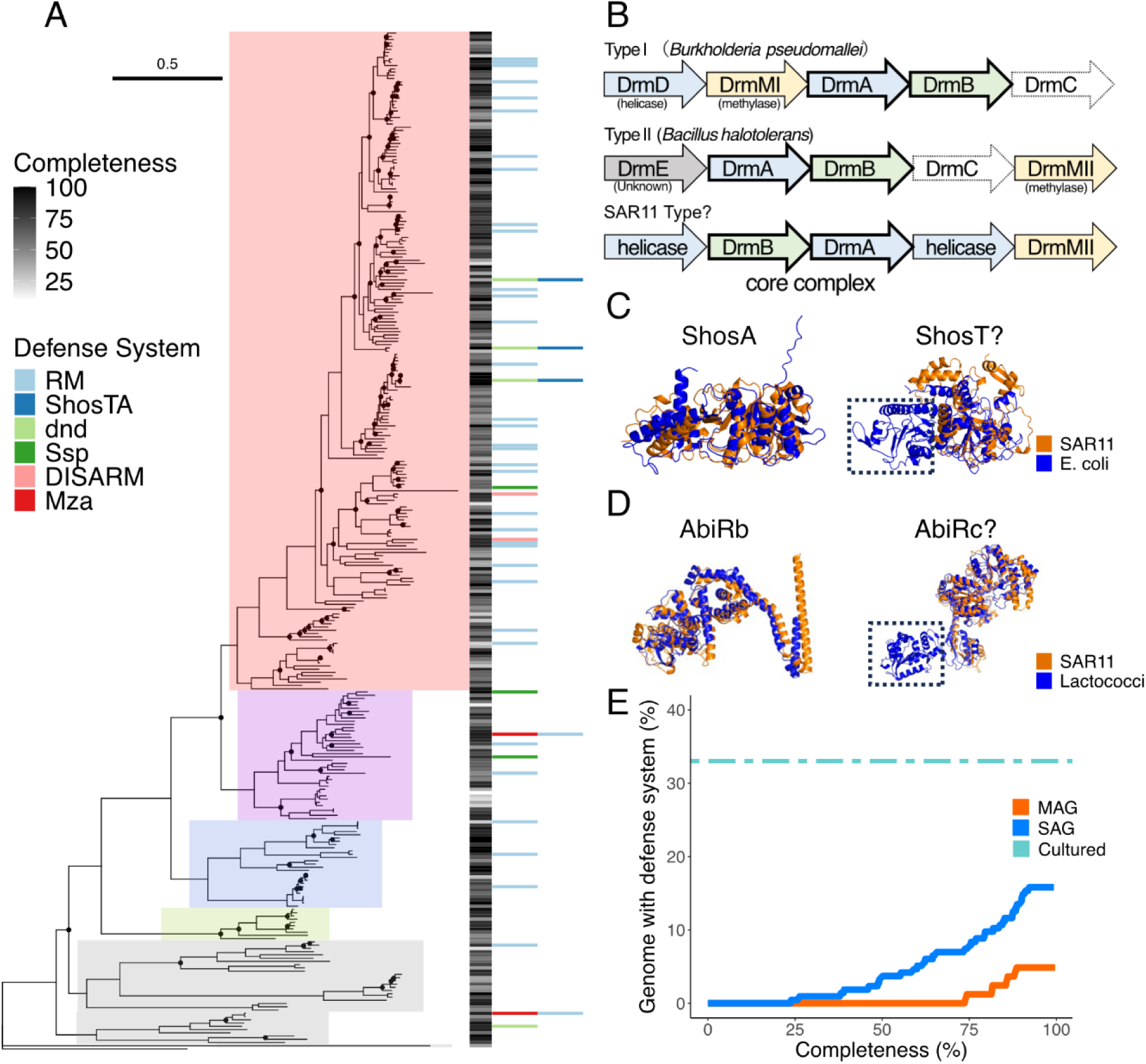
Defense genes identified in SAR11 genomes. **(A)** Sporadic distribution of defense systems. Heatmap colors indicate genome completeness and presence of defense systems. **(B)** Operon structure of known DISARM systems and AG-447-N21 coding DISARM-like system. The colors indicate function as follows: helicase (blue), methylase (yellow), DUF1998-containing protein (green), gray (unknown), and PLD domain (white; not required for defense). **(C)** Predicted structure comparison of the known ShosTA proteins from DefenseFinder wiki (blue) and the ShosTA-like proteins in HTCC1002 (orange). Domain loss is highlighted by the dotted box. **(D)** Predicted structure comparison of known AbiRbc proteins from DefenseFinder wiki (blue) and AbiRbc-like proteins in AG-313-A04 (orange). **(E)** The cumulative proportion of genomes with detected viral defense systems is plotted against genome completeness. The cumulative proportion at each completeness value represents the fraction of genomes at or below that completeness in which at least one defense system (from panel A) was detected. Solid blue and red lines show the cumulative proportion for SAGs and MAGs, respectively. The light blue dashed line indicates the detection rate in 24 cultured genomes, serving as a reference.

A previous study showed that only a few deep sea SAR11 genomes possess part of the CRISPR-Cas-like gene set (93). As in the previous study, Cas-like genes without CRISPR array were detected in two deep sea SAGs (AG-337-I02 and AG-410-B18). Three cultured-strains genomes and one MAG possess *dnd*, a dsDNA phosphorothioation (PT)-based defense system (94). In addition, we identified Ssp-like operons on two SAR11 genomes **(Fig. 4A)**. Same as the *dnd*, Ssp is a PT-dependent system, modifing DNA by SspBCD and cleaving foreign ssDNA by SspE (95,96). The SAR11 Ssp-like operons encode proteins structurally similar to the known SspBCD and an uncharacterized protein shorter than the known SspE. Although experimental validation is needed, this finding indicates that PT-based modifications are used to distinguish self from foreign DNA in the SAR11 clade.

We also identified Mza, DISARM, and ShosTA-like operons **(Fig. 4A)**. Mza defends against ssDNA phages, although its mechanism remains unknown (97). The DISARM-like operon in SAR11 encodes DrmAB, DrMII-type methyltransferase, and helicases **(Fig. 4B)**, whereas DrmC, nonessential for defense, appears absent (98). The distinct gene order suggests a divergent DISARM-like configuration. The ShosTA toxin-antitoxin system was detected in three SAR11 genomes. The SAR11 ShosT protein is approximately 100 amino acids shorter than the *E. coli* ShosT but retains the phosphoribosyl transferase domain **(Fig. 4C)**. This suggests that the SAR11 clade may have acquired a truncated but functional toxin, likely through genome reduction. AbiRb-and AbiRc-like genes were also present as operons, but the AbiRc-like gene in SAR11 lacks its terminal domain **(Fig. 4D)**. Moreover, AbiRa is a sigma70 factor, yet the only sigma70-type RNA polymerase subunit is present in the SAR11 genome, it is not located within this operon. Therefore, it remains unclear whether this AbiR-like genes functions as a known abortive infection system, and we did not include it as a defense system found in this study.

Previous limited identifications of viral defense genes in SAR11 may be partly due to the presence of these overlooked putative defense systems that slightly differ from experimentally validated defense system components (**Fig. 4BCD**). When we focused on high-quality genomes, defense systems were found in at least one-third of the cultured strains and one-quarter of the SAGs. In contrast, defense systems were rarely detected from the MAGs **(Fig. 4E)**. This result suggests that the SAR11 clade harbors more diverse and abundant defense systems than previously inferred from the MAG-based analyses (20).

## Discussion

As genomic data continue to expand exponentially, functionally unknown genes are becoming more prevalent, and their study—termed Functional Unknomics (26)—is gaining attention. Previous research on genes of unknown function has primarily focused on model organisms (83,26,99) or large-scale sequence analyses of whole ecosystems, involving MAGs and SAGs (25,29,30). This study represents the first systematic investigation focused on genes of unknown function in the SAR11 clade as one of the non-model organism lineages.

In the SAR11 clade, one of the most abundant marine bacterial lineages, a large fraction of OGs remains functionally uncharacterized despite their crucial roles in marine microbial ecology. To address this issue, we clustered 306,416 protein-coding genes in 321 SAR11 genomes into 5,603 OGs and classified 3,255 of them (58.1%) as unknome based on the COG database **(Fig. 1CD)**. Utilizing genome context and predicted protein structures, we successfully obtained functional information for 69 out of the 108 core unknomes **(Table S5 and Supplemental Text).** These included many transporter genes, supporting the strategy of SAR11 that specializes in substrate uptake, as proposed in previous studies (3,9,10). Furthermore, our analysis confirmed that the utilization of genomic context and predicted protein structures is valuable even for non-model organisms. Especially, structural similarity search proved particularly effective in functional annotation (**Fig. 2D, S7, S8 and Supplemental Text**). This may reflect both the accumulation of neutral mutations during the divergence of SAR11 clade from well-studied model organisms and shifts in genomic properties, including reduced GC content, leading to the presence of many remote homologs (5). Although structural similarity search can find distant relationships between SAR11 proteins and those in the PDB, the latter also lack functional information, such as the DUF2237-containing protein. In this study, we addressed this problem by manually examining the individual results. This involved cross-referencing them with databases such as PDB, Pfam, COG, KO, Uniprot, and STRING, and conducting literature searches for any reported functional insights **(Supplemental Text)**. This careful curation allowed us to focus on previously uncharacterized genes potentially support ecological success of SAR11 clade (**Supplementary Text**).

We found functionally unknown genes specifically related to the marine environment, which are probably crucial for the ecology of the SAR11 clade **(Fig. 3A)**. Among them, we found the DUF2237-containing proteins, which were previously thought to be related to proteorhodopsin (87). The predicted protein structure and docking simulation results suggested that DUF2237 is likely to be an enzyme associated with purine nucleotide-containing compounds **(Fig. 3CDE)**. This result may indicate that the DUF2237 protein binds to second messengers, such as c-di-GMP and cAMP, and is thus involved in phototaxis. Alternatively, it might interact with NADH, indicating involvement in redox processes. This finding collectively suggests that DUF2237 participates in fundamental cellular processes, extending beyond a direct link to phototaxis. Although definitive conclusions require further experimental validation, the possible binding of DUF2237 to these compounds suggests that it represents a novel nucleotide-binding motif and it may play a key role in marine bacterial ecosystems.

We explored the unknome genes to identify viral defense systems that may offer insights into top-down factors contributing to ecological success of the SAR11 clade. Previous studies suggested that SAR11 possesses only limited viral defense capacity (3,20). However, our comprehensive genomic analysis revealed a scattered distribution of diverse viral defense systems **(Fig. 4A)**. This pattern is consistent with the high intraspecific recombination rates of SAR11 genes (22). While the previous hypothesis proposed that a slow growth rate of SAR11 made it less vulnerable to phage lysis (15), the discovery of antiviral defense systems indicates that SAR11 face viral pressure, requiring more active resistance to viral infection than previously appreciated. The recent discovery of anti-RM systems in the Pelagiphage genome further supports the existence of the arms race between *Ca.* Pelagibacter and the Pelagiphage (80). The presence of defense systems might potentially explain the unique ribosome-deprived phenomenon known as ‘zombie cells’ in SAR11 upon viral infection, possibly resulting from abortive infection (20). Moreover, the discovery of the Ssp and Mza systems in the SAR11 genomes, both of which are known to target ssDNA viruses, suggests the potential existence of undiscovered ssDNA Pelagiphages. Given the high intraspecific recombination rate of SAR11 genes (22), the sporadic distribution of defense systems likely reflects population-level sharing of this repertoire, rather than frequent transfer across distantly related taxa. Such population-level sharing may help SAR11 balance the trade-off between genome streamlining and antiviral defense. This interpretation is consistent with the concept of a bacterial pan-immune system, in which defense genes are frequently gained and lost among closely related strains (100), and also consistent with the KoM hypothesis, which proposes SAR11 maintains its dominance through high diversity and efficient horizontal gene transfer driven by high contact frequency of closely related strains (16,21). Collectively, these observations suggest a complex evolutionary interplay between SAR11 and its phages, likely shaped by genome streamlining, horizontal gene transfer, and diverse defense strategies.

Our results also underscore the importance of genome reconstruction methods in evaluating the distribution of viral defense systems. Compared with SAGs and genomes from cultured strains, viral defense genes were underrepresented in MAGs **(Fig. 4D)**, which may have led to the previous observation that SAR11 possesses little viral defense capacity (3,20). This underrepresentation likely stems from limitations in MAG assembly and binning, reflecting the high variability of specific genomic regions (e.g., hyper variable regions and metagenomic island) (6,23,24). Consequently, SAG analysis provides a powerful approach to capture diversity of defense systems, as it circumvents the challenges associated with MAG construction (37).

Looking forward, further advances in automated gene function annotation will be essential to keep pace with the rapid expansion of microbial genomic data and to complement meticulous but low-throughput manual annotation. By developing robust analytical approaches and moving beyond model organisms, we can uncover novel biological and ecological functions of microbes, providing deeper insight into the diversity and ecological roles of microbial life in natural environments. Bioinformatics-driven Functional Unknomics is poised to become a major frontier in microbial ecology, particularly for lineages that remain difficult to culture.

## Acknowledgments

The authors would like to acknowledge Wataru Iwasaki for providing computational resources. Computation time was provided by the SuperComputer System, Institute for Chemical Research, Kyoto University.

## Author contributions

K.T., S.N., Y.N., and K.O. conceived and designed this study. S.N. collected data and performed the bioinformatic analysis and data visualization throughout this study. K.T. performed the meta-transcriptomic read mapping analysis. K.T., Y.N., K.O., T.D., K.H., and S.Y. provided critical interpretations of the data. S.N. drafted the manuscript. All authors revised and approved the final version.

## Conflicts of interest

The authors declare that there are no conflicts of interest.

## Funding

This work was supported by JST GteX Grant Number JPMJGX23B2 and JSPS KAKENHI Grant Number JP22H04925 and JP24KJ0979.

## Data availability

The code and datasets generated and analyzed in this study are available in our Zenodo repository, https://doi.org/10.5281/zenodo.16318636.

## Notes

https://doi.org/10.5281/zenodo.16318636

### Competing Interest Statement

The authors have declared no competing interest.

### Summary of Updates

Error correction in the author suffix.

