## Supplemental Figures for "Functional Unknomics of the SAR11 clade using bioinformatics approaches"

**Supplemental Figures of "Functional Unknownmics of the SAR11 clade using bioinformatics approaches"**

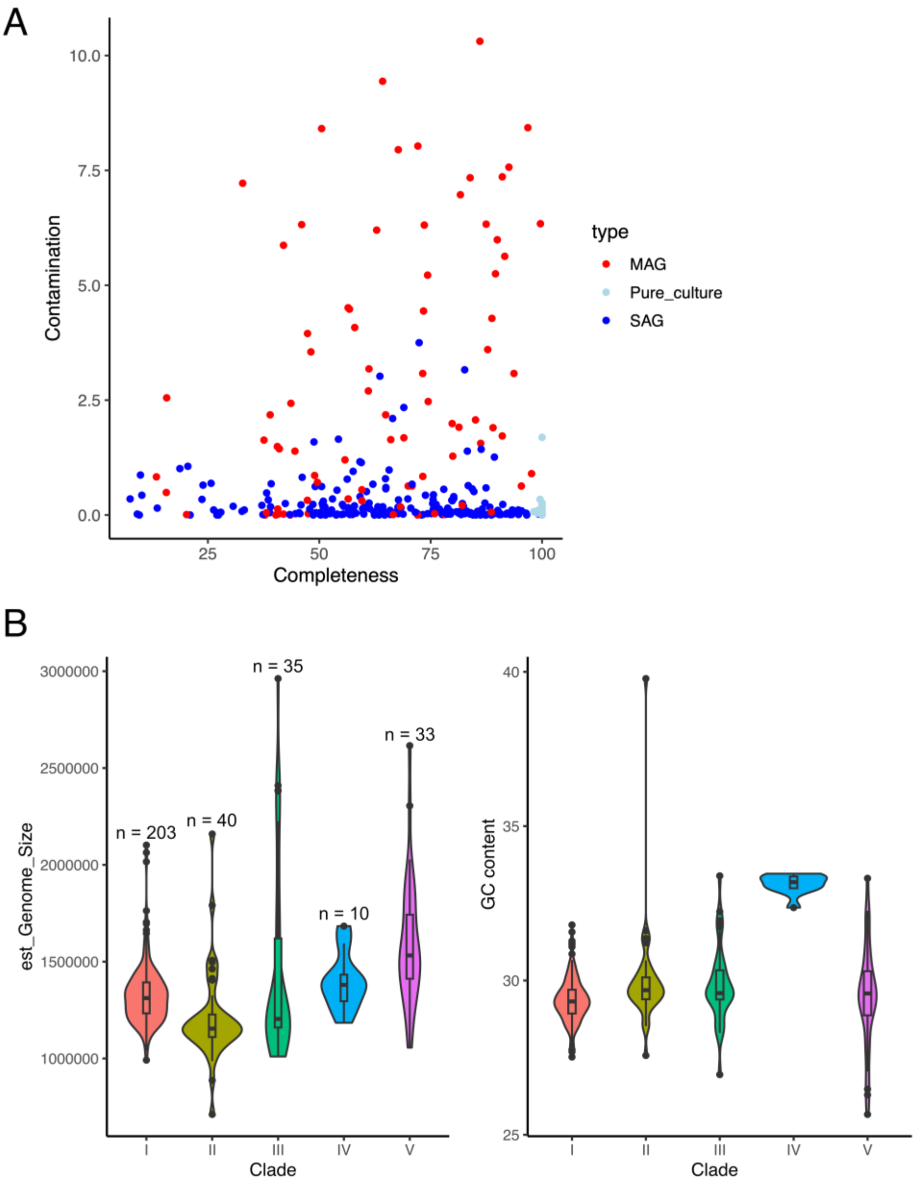


**Supplement Figure 1. Quality and sequence properties of genomes used in this study.**

**(A)** Scatter plot of the Completeness and Contamination scores calculated by CheckM2. The colors indicate genome types as follows: Metagenome-assembled genomes (red), Cultured strain (light blue), and Single-amplified genomes (blue). **(B)** The estimated genome size and GC contents of the five clades.

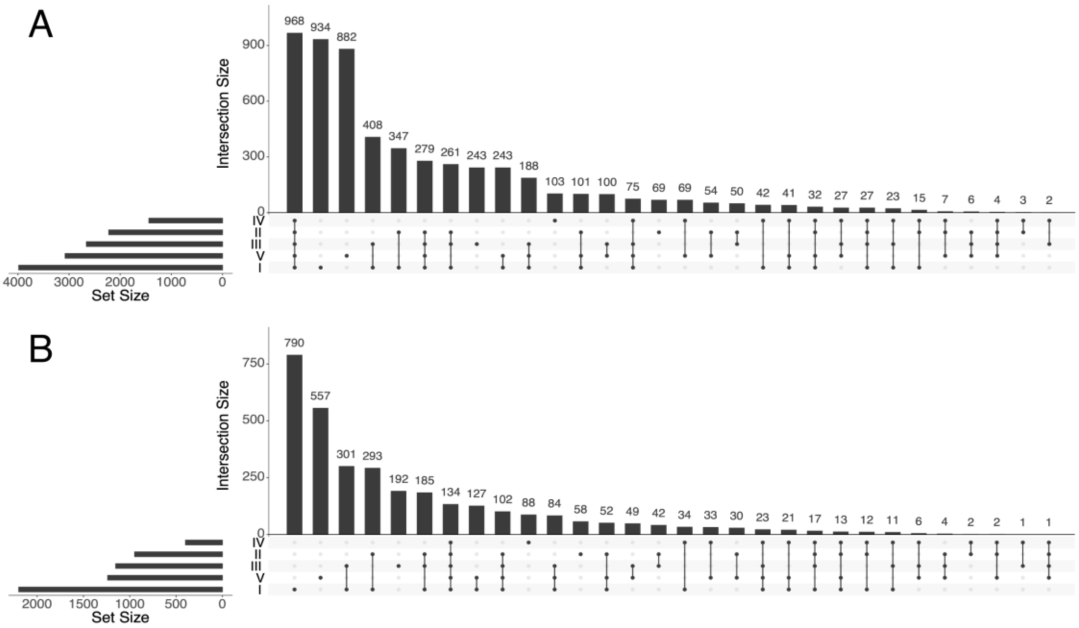


**Supplement Figure 2. Distribution of OGs among the five SAR11 subclades**

**(A)** Upset plot of all OGs distribution among the five SAR11 subclades. **(B)** Upset plot of the unknome OGs distribution among the five SAR11 subclades.

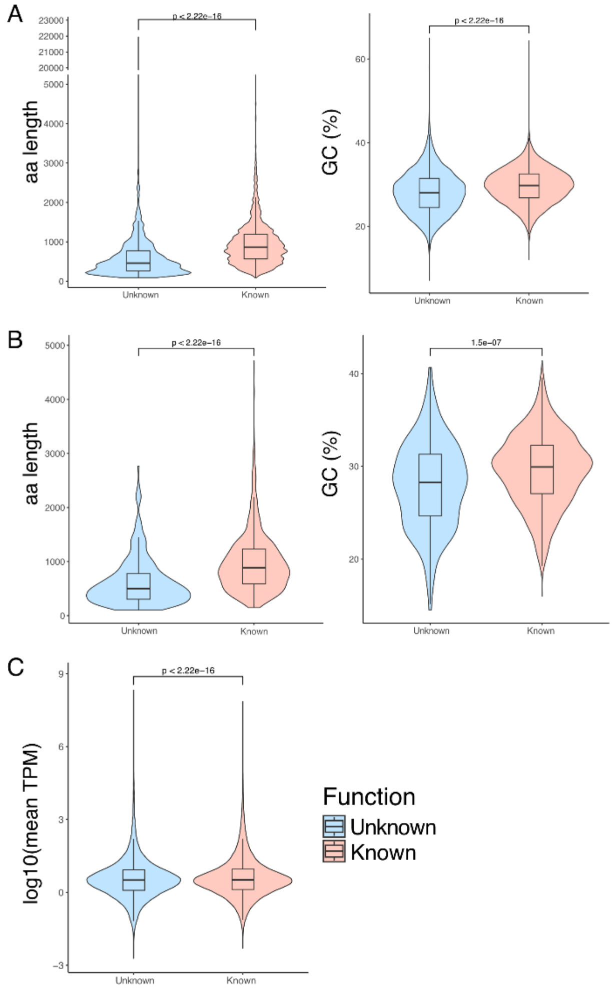


**Supplement Figure 3. Comparison of size and GC content between functionally known and unknown genes in SAR11 clade.**
**(A)** The amino acid length and GC content distribution of all SAR11 genes analyzed in this study. **(B)** The amino acid length (< 5,000 aa proteins) and GC content distribution of *Ca.* P. *ubique* HTCC1062 strain. **(C)** The average transcript per million (TPM) value of each gene calculated from the read mapping of Tara Oceans meta-transcriptome data to the SAR11 genomes analyzed in this study.

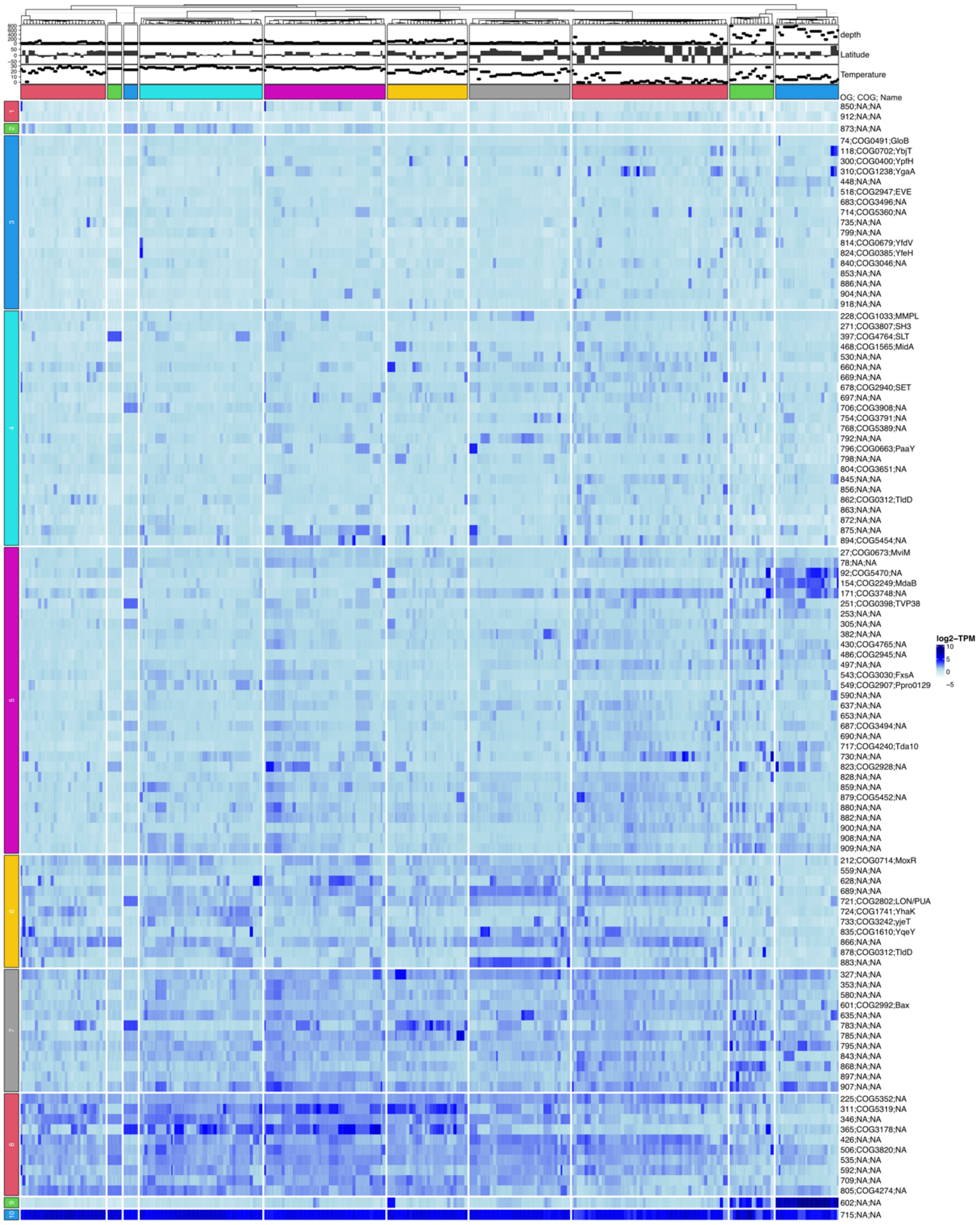


**Supplement Figure 4. Meta-transcriptome expression profiles of the 108 core unknome OGs.**

The heatmap shows expression profiles of each core unknome OGs based on TPM value. The heatmap was clustered using the default settings of ComplexHeatmap in R (n = 10, Euclidean distance and complete linkage). Plots above the heatmap indicate the depth, latitude, and temperature for each sampling site. The vertical axis represents each core unknome OG and assigned COG information.

A

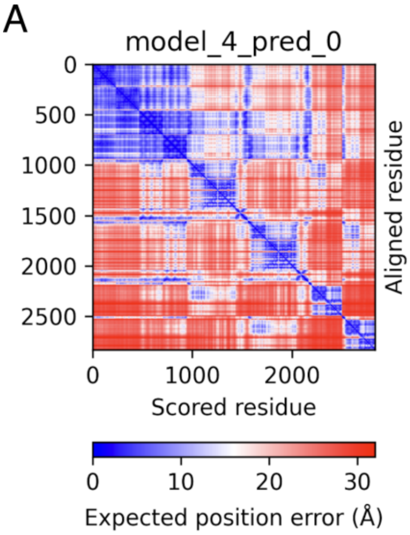


B

| Chain | color | pDockQ_i |
|---|---|---|
| A (OG688) | pink | 0.498444 |
| B (OG688) | pink | 0.509799 |
| C (OG1772) | orange | 0.529385 |
| D (OG1772) | orange | 0.523811 |
| E (OG1815) | yellow | 0.268428 |
| F (OG1815) | yellow | 0.274739 |
| G (OG1814) | gray | 0.363983 |
| H (OG1814) | gray | 0.336381 |

**Supplement Figure 5. Predicted aligned error plot and pDockQ2 scores of the putative transporter complex in Figure 2B.**

**(A)** A predicted aligned error plot of the top-ranked predicted structure model. **(B)** pDockQ2 score of each chain. The minimum pDockQi score is 0.268 (chain E), indicating the potential for complex formation.

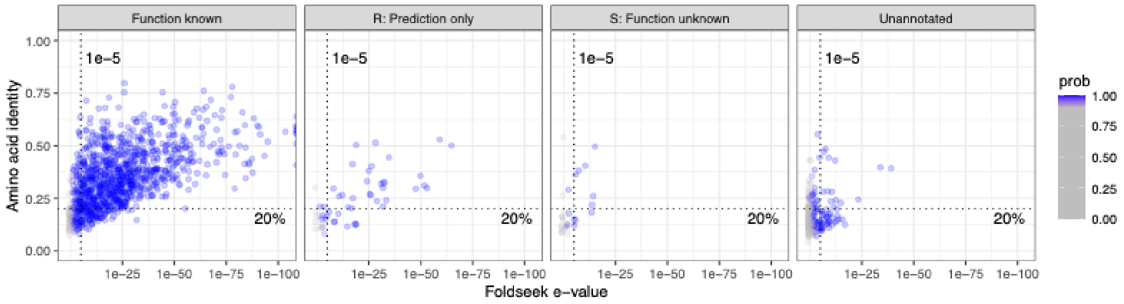


**Supplement Figure 6. Results of protein similarity search in IMCC9063 strain.** Scatter plots of all IMCC9063 strain proteins compared with known protein structures in the Protein Data Bank. The dot lines show e-value = 1e-5 and amino acid identity = 20%, highlighting potential remote homologs.

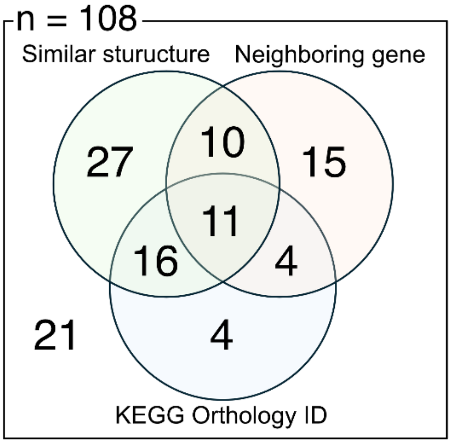


**Supplement Figure 7. Overlap of functional annotations obtained for the core unknome OGs by three distinct approaches.**
A Venn diagram shows the annotation results for 108 core unknome OGs in the SAR11 clade. Each circle represents the number of OGs that received a hit from a specific annotation approach: predicted structure similarity search to PDB (green), neighborhood gene analysis (orange), or KEGG Orthology (KO) ID assignment (blue). Detailed functional investigations of each OG are provided in the Supplemental Text.

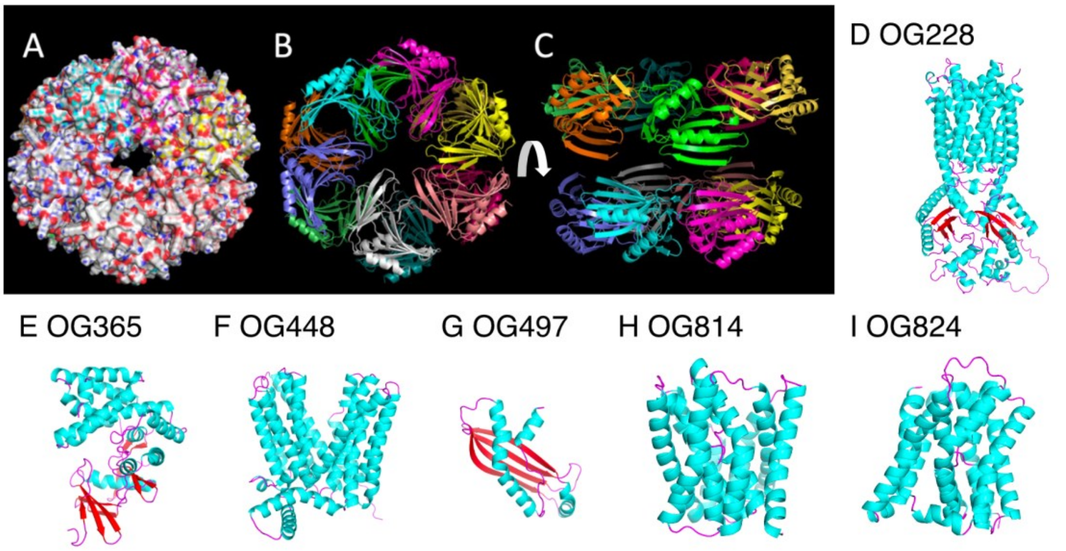


**Supplement Figure 8. Predicted unknome protein structures of putative transporter, channel, or metabolic enzyme.**

(A-C) Predicted homo-12mer structure of OG305, putative channel protein, from *Ca*. P. ubique HTCC1062 strain. Each color shows each chain. (D-I) Predicted monomer structures of the unknomes from *Ca*. P. ubique HTCC1062 strain. Colors indicate secondary structure elements: alpha-helices (light blue), beta-sheets (red), and loops (pink). (D) Putative transporter. (E) Putative AmpK-like anomeric sugar kinase. (F) Putative AmpG-like transporter. (G) Putative translocation-associated chaperone. (H) Putative sodium/solute symporter. (I) Putative bile acid/sodium symporter.

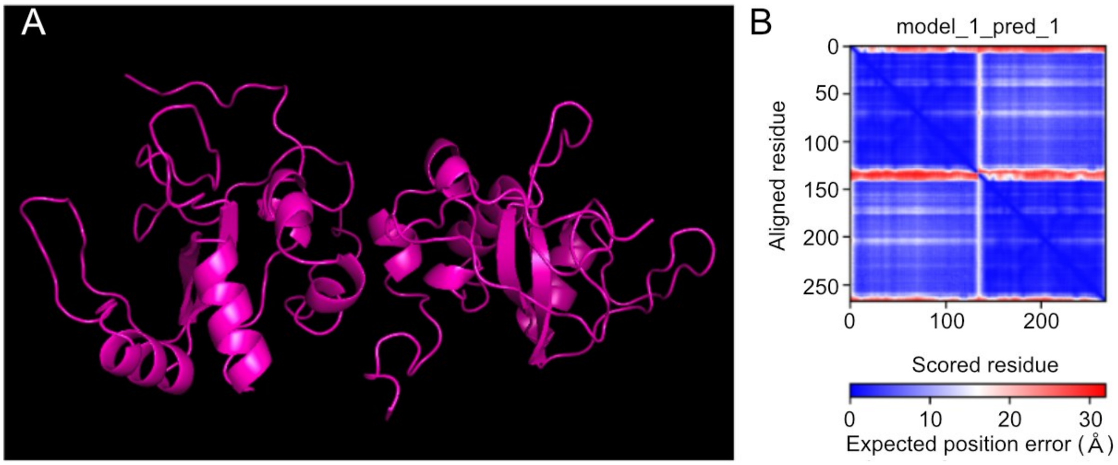


**Supplement Figure 9. Predicted DUF2237 homodimer structure.**
**(A)** Predicted homo-dimer structure of DUF2237 protein on *Ca.* P. ubique HTCC1062 strain by AlphaFold2 multimer. **(B)** A predicted aligned error plot of the predicted homo-dimer structure of the DUF2237 protein.

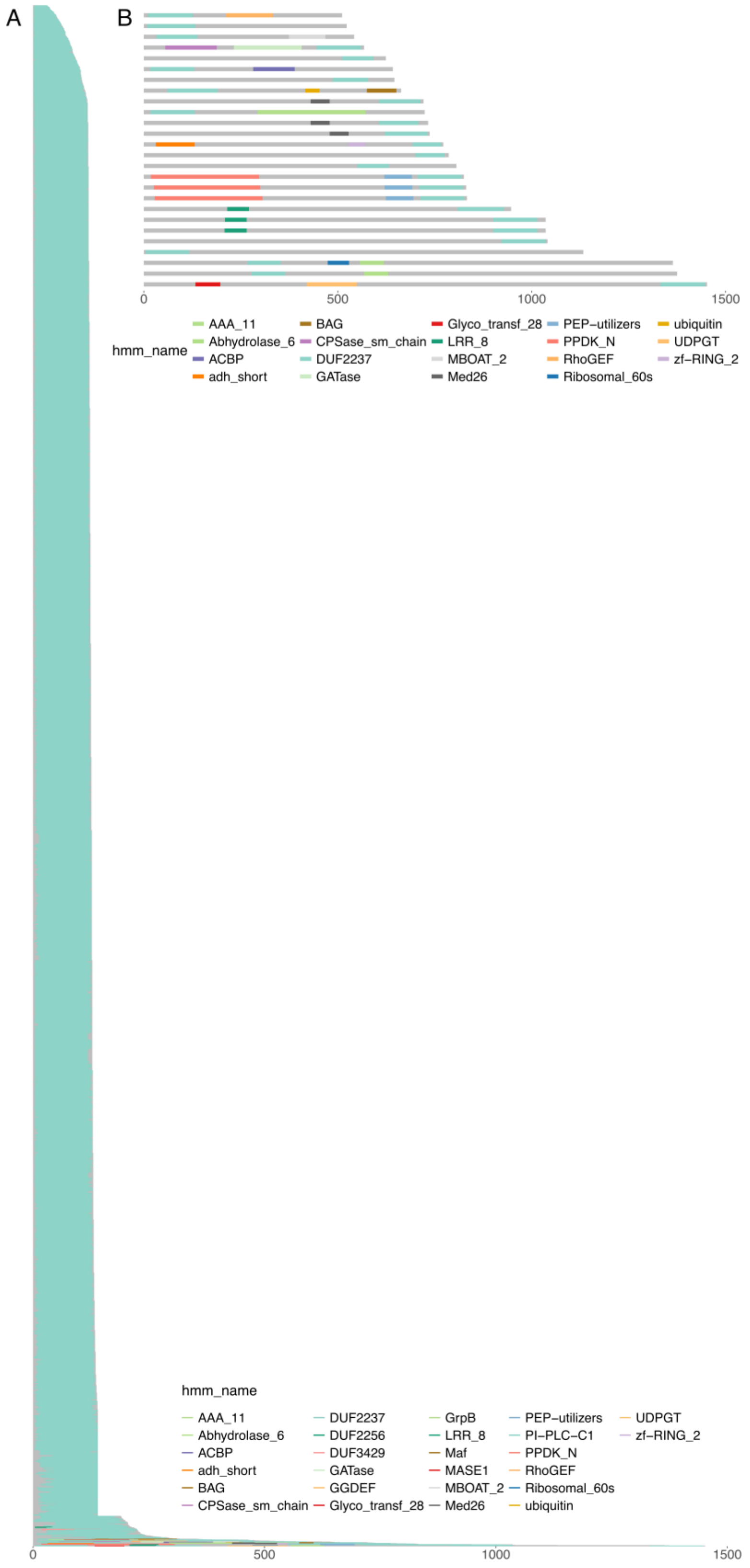

A
B
0
500
1000
1500
hmm_name
AAA_11
Abhydrolase_6
ACBP
adh_short
BAG
CPSase_sm_chain
DUF2237
GATase
Glyco_transf_28
LRR_8
MBOAT_2
Med26
PEP−utilizers
PPDK_N
RhoGEF
Ribosomal_60s
ubiquitin
UDPGT
zf−RING_2
DUF2256
DUF3429
GGDEF
GrpB
Maf
MASE1
PI−PLC−C1

**Supplement Figure 10. Domain architecture of 7,963 DUF2237-containing proteins in the Pfam database.**

**(A)** Domain architecture of 7,963 DUF2237-containing amino acid sequences in the Pfam database. The x-axis represents the length of the amino acid sequences, while the y-axis arranges the sequences in order of size. The colors indicate the presence of protein domains within each sequence. **(B)** Domain compositions of over 500 aa DUF2237-containing proteins in panel A.

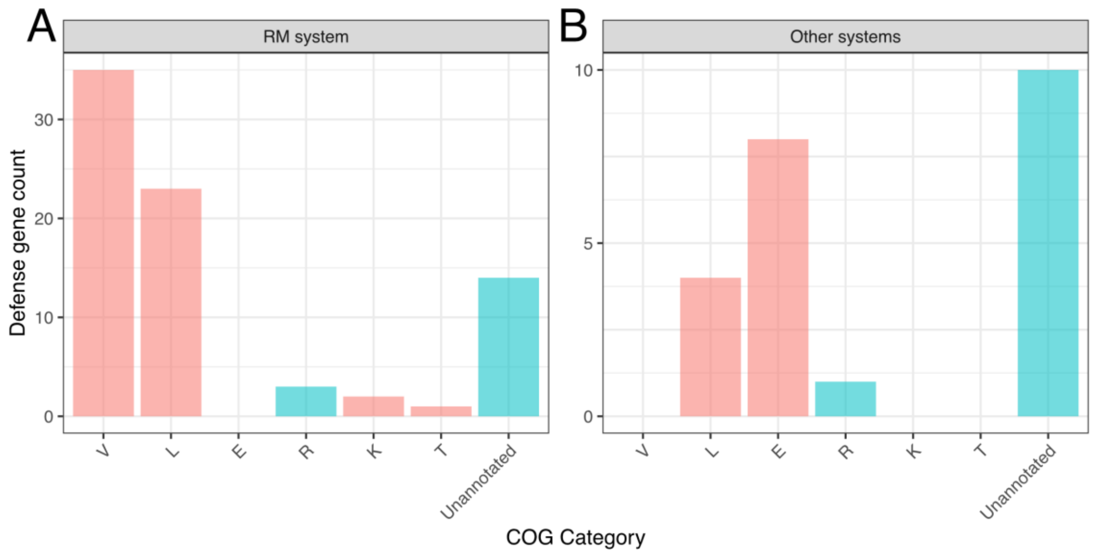


**Supplement Figure 11. COG classification result of DefenseFinder-detected antiviral defense genes.**

**(A)** Bar plot showing the number of genes in each COG category among all DefenseFinder-detected restriction-modification (RM) system genes. The unannotated represents genes without a COG annotation. **(B)** Bar plot showing DefenseFinder-detected defense genes excluding RM systems.

**Supplement Tables**

Table S1. SAR11 genomes used in this study.

Table S2. List of ENA_Run_IDs used for the meta-transcriptomic read mapping.

Table S3. Docking simulation results and compounds list.

Table S4. Sequence similarity-based ortholog annotation results.

Table S5. Function prediction summary of the 108 core unknome OGs.

Table S6. SAR11 defense systems found in this study.

**Supplement Text**

Annotation results for the 108 core unknome OGs.
