## Supplemental Text for "Functional Unknomics of the SAR11 clade using bioinformatics approaches"

### Supplemental manuscript for "Functional Unknomics of the SAR11 clade using bioinformatics approaches"


This manuscript contains annotation results for the 108 core unknome.

#### 1. Functions could be estimated from combined structural similarity search, neighboring genes, and literature information (8 OGs)

**OG27 MviM**

OG27 was annotated as COG0673 and identified as the gene known as *mviM*. There are four previous studies on *MviM*, all of which report its inclusion in sugar catabolic operons and its involvement in redox reactions targeting specific hydroxyl groups of sugars (1–4). However, types of sugars and the neighboring sugar metabolism genes vary among these studies. In the case of SAR11, the ligand could not be inferred from the genome context. Structural predictions revealed that MviM possesses a Rossmann fold, a nucleotide-binding domain, confirming that it is an NADH-dependent oxidoreductase (5).

**OG300 PE8 esterase**

OG300 was annotated as K06999 and COG0400, corresponding to the gene known as ypfH in *Escherichia coli*. YpfH is known to exhibit palmitoyl-CoA esterase activity. Structural similarity analysis revealed that OG300 shares similarities with multiple esterases, including those with PDB entries 5DWD, 4FHZ, and 5SYN, suggesting that it is likely a homolog of the PE8 esterase. Notably, the PE8 esterase from *Pelagibacterium halotolerans*, which shows high structural and sequence similarity, has been reported to hydrolyze the prochiral compound dimethyl 3-(4-fluorophenyl) glutarate (6–8). In the genomic neighborhood, a 5-methylcytosine-specific restriction enzyme gene (mcrA), known to be involved in defense against viral infection (9), was conserved, although its functional relationship with OG300 remains unclear.

**OG353 Tol-Pal system inner membrane component**

OG353 was initially classified as a functionally unknown membrane protein. Structural similarity analysis revealed that it shares a similar fold with TolA, a component of the Tol-Pal system (e.g., PDB entries 3QDR, 1S62). In its genomic neighborhood, an operon containing genes related to the Tol-Pal system, including tolB, tolQ, and ompA, was conserved. These findings suggest that OG353 is likely a remote homolog of TolA.

**OG549 Putative light-related oxidoreductase**

OG549 was annotated as COG2907, a gene of previously unknown function. However, structural predictions revealed that it is an $NAD^+$/$FAD^+$-binding oxidoreductase. In the gamma-proteobacterium *Acinetobacter baumannii*, transcription of the operon containing this gene has been shown to increase under blue light exposure (10). In Aeromonas salmonicida, another gammaproteobacterium, transcription of the gene decreases when cultured in riboflavin-supplemented medium, suggesting a link to fatty acid metabolism (11). A similar operon structure, including OG683, is also observed in SAR11. However, previous studies culturing SAR11 under light conditions did not detect any significant increase in transcription (12).

**OG683 Putative pigment synthesis gene**

OG683 was annotated as K09701 and COG3496, and contains a DUF1365 domain, classifying it as a domain of unknown function. In the gammaproteobacterium *A. baumannii*, transcription of this gene is known to increase under blue light exposure (10). In a previous study on *A. salmonicida*, transcription levels decreased when cultured in riboflavin-supplemented medium, suggesting a potential role in fatty acid metabolism (11). Additionally, a possible association with the carotenoid lutein has been suggested in Cucurbita (pumpkin) (13). In contrast, no significant increase in transcription was observed in SAR11 when cultured under light conditions (12). Structural similarity analysis revealed that OG683 shares a similar fold with the fatty acid-binding protein Ycf58 (PDB 3BDR). Taken together, these findings suggest that OG683 may be involved in pigment biosynthesis related to light utilization.

**OG866 Membrane anker protein AprM**

OG866 was initially classified as a functionally unknown membrane protein. However, the presence of neighboring genes encoding adenylylsulfate reductase subunit beta (aprB) and subunit alpha (aprA) suggested that OG866 may be aprM—the membrane anchor of the adenosine-5′-phosphosulfate reductase (AprMAB) complex, which catalyzes the oxidation of sulfite to sulfate (14). To test this hypothesis, we performed a complex structure prediction using the amino acid sequences of these three genes by AlphaFold-multimer, which successfully yielded a predicted complex. No known proteins with a similar structure were found in the PDB. This complex is known to catalyze the oxidation of sulfite to sulfate (15).

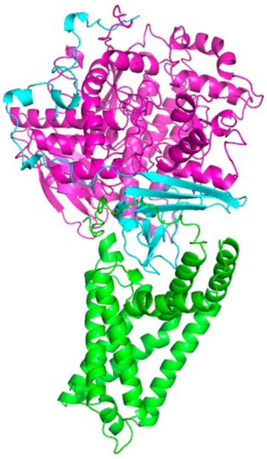

Predicted complex structure of OG866 and its two downstream genes in *Ca.* P. ubique HTCC1062 strain: OG866 (green), AprA (pink), and AprB (light blue).

**OG882 Exonuclease VII small subunit protein**

Sixteen out of 173 sequences showed sequence similarity to the Exonuclease VII small subunit in the COG database (maximum identity: 34.2%), and one sequence was annotated as COG4942 (EnvC; identity: 29.4%). The predicted compact hairpin-like helical structure of OG882 is also consistent with that of the Exonuclease VII small subunit, supporting the possibility that OG882 represents a homolog of this protein. This hypothesis is further supported by annotations in the STRING database (e.g., AAZ21430.1), where some of these sequences are also predicted to encode Exonuclease VII small subunits.

**OG909 Cell division protein FtsL**

In SAR11 genomes, the neighboring gene OG169 (COG0769; UDP-N-acetylmuramoyl tripeptide synthase) is conserved, suggesting a potential link to cell envelope biosynthesis. Among the 163 sequences classified under OG909, four showed significant similarity to COG4839 (maximum amino acid identity: 31.3%), suggesting that OG909 may be a homolog of FtsL2, a cell division protein FtsL. However, similarities were also observed with other COGs, including COG4942 (30.3% identity), COG5462 (31.8% identity), and COG1593 (21.3% identity), making it difficult to predict the function of OG909 based solely on database annotation. A structural similarity search using the predicted 3D structure of OG909 from the *Ca.* P. ubique HTCC1062 strain revealed structural similarity to the well-characterized *E. coli* FtsL protein (16) (Foldseek probability = 0.98), although the amino acid sequence identity remained low at 18.8% (see figure below). These results suggest that OG909 may represent a remote homolog of FtsL. These results were further supported by neighboring gene information from the STRING database.

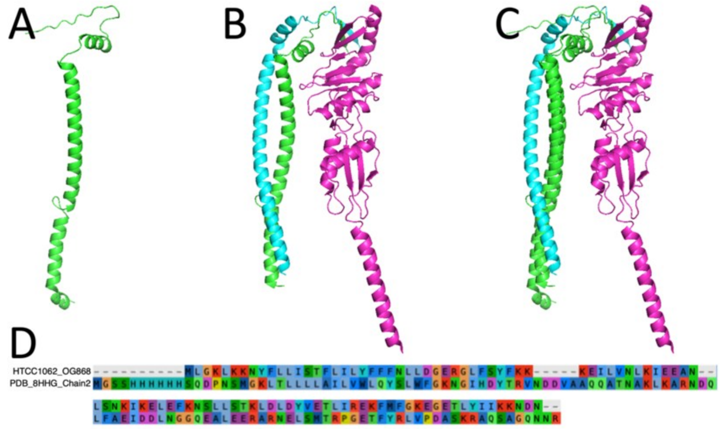


**OG868 and known FtsB** A. Predicted structure of OG868. B. Structure of the *E. coli* K-12 FtsBLQ complex (PDB: 8HHG). C. Structural superposition of A and B (Foldseek probability = 0.98, sequence identity = 18.8%; PyMOL RMSD by super = 5.869 Å). D. Amino acid sequence alignment between OG868 from *Ca*. P. ubique HTCC1062 and FtsB (PDB 8HHG, Chain 2).

#### 2. Functions are known from literature information (7 OGs)

**OG310 DedA family flippase PetA（YgaA）**

OG310 was annotated as COG1238 and identified as a membrane protein of the DedA superfamily, which is commonly found in Gram-negative bacteria. In March 2023, it was suggested that this protein interacts with the phospholipid-bridging protein AsmA to mediate phospholipid flipping (17). Subsequently, in May 2023, its function as a phosphatidylethanolamine transporter was experimentally confirmed in *Bacillus subtilis*, and the protein was named PetA (18).

**OG506 Transcriptional regulators including DUF1013, TrcR**

OG506 encodes a protein containing the DUF1013 domain, a domain of unknown function. In SAR11 genomes, two neighboring genes—OG507 and OG442—encode the non-essential large subunit ribosomal protein L31 and the Elongation Factor P, respectively, suggesting that the functionally uncharacterized OG506 may be related to translation. A previous study screening cell cycle mutants of the model alpha-proteobacterium *Caulobacter crescentus* identified a DUF1013-containing protein that binds RNA polymerase and affects both transcription initiation and elongation, and this protein was named TrcR (19). In bacteria, it is well established that transcription elongation by RNA polymerase II and translation by ribosomes are spatially and temporally coupled (20,21), raising the possibility that OG506 may be involved in this mechanism.

**OG518 RNA binding protein**

OG518 was annotated as COG2947 and identified as a protein of unknown function containing an EVE domain. Structural similarity analysis revealed that it shares structural features with RNA-binding proteins, such as the one represented by PDB entry 2GBS. The EVE domain is known to function as an RNA-binding motif (22). Two evolutionary types of EVE domains have been described—one evolving rapidly and the other more slowly—and the latter is found in alpha-proteobacteria, including the SAR11 clade (23). Proteins containing the alphaproteobacterial-type EVE domain are suggested to be involved in translation, respiration, and cytochrome c binding through recognition of 5-methylcytosine modifications in tRNA (23).

**OG637 Phospholipid bridge to outer membrane**

OG637 was identified as a gene of unknown function containing the DUF3971 domain. Recent studies have suggested that its *E. coli* homolog, YhdP, is involved in maintaining outer membrane phospholipid homeostasis (24,25). In October 2023, a preprint was published that elucidated its molecular mechanism, reporting that YhdP functions as a phospholipid bridge to the outer membrane (26).

**OG785 BamC homolog of Alphaproteobacteria, BamF (DUF3035)**

OG785 encodes a protein containing a DUF3035 domain, and its genomic neighborhood includes OG490 (COG0742; 16S rRNA G966 N2-methylase RsmD). DUF3035 is an uncharacterized domain found exclusively in Alphaproteobacteria and is known as BamF. BamF is a lipoprotein component of the outer membrane β-barrel assembly machinery (BAM) complex (27).

**OG845 DCC family thioredoxin**

OG845 was identified as a gene of unknown function containing the DUF393 domain. DUF393 is known to share structural similarity with thiol-disulfide oxidoreductases (28), suggesting that OG845 also functions as a thioredoxin.

**OG878 Metalloprotease TldD**

OG878 was annotated as K03568 and predicted to be a metalloprotease belonging to the TldD family (29). Structural similarity analysis confirmed that it shares a fold with known TldD proteins. TldD is known to form a heterodimer with the TldE protein and function as a metalloprotease (30). In the SAR11 genomes, functionally uncharacterized genes corresponding to TldE (OG862) were also identified, suggesting the possibility that OG878 forms a heterodimer with it.

### 3. Function predicted by predicted structure (36 OGs)

**OG78 Methyltransferase**

OG78 is an uncharacterized gene containing a methyltransferase domain. Structural similarity analysis revealed resemblance to multiple methyltransferase proteins, including tRNA methyltransferase (PDB: 8K1F).

**OG92 DUF1330-containing cofactor-independent oxygenase**

OG92 harbors a domain of unknown function, DUF1330. The DUF1330 is primarily found in Proteobacteria and Actinobacteria and has been shown to be essential for the biosynthesis of L-allo-isoleucine, a precursor of coronatine, in Pseudomonas (31). A structural similarity search revealed three experimentally determined structures (PDB: 2FIU, 3LO3, 3DCA), all of which remain functionally unannotated. These structures are all deposited as homodimers. Notably, the structural features resemble those of cofactor-independent monooxygenases such as SnoaB and LsrG (32), suggesting that OG92 may function as an oxygenase capable of utilizing molecular oxygen directly.

**OG118 DUF2867-containing redox enzyme associated with the NADH:ubiquinone oxidoreductase complex**

OG118 is an uncharacterized gene also known in *Escherichia coli* as YbjT (COG0702). This protein harbors the DUF2867 domain. Structural prediction revealed the presence of a nucleotide-binding motif consistent with a Rossmann fold (5), and no transmembrane helices were predicted. A structural similarity search indicated that OG118 shares significant structural homology with the NADH-binding proteins of the eukaryotic respiratory complex I. Furthermore, co-expression data from the STRING database suggest a potential association with bacterial complex I gene clusters (NuoABCDGIM). Collectively, these findings suggest that OG118 may function as a redox enzyme associated with the NADH:ubiquinone oxidoreductase complex.

**OG253 Fe(II)/2-oxoglutarate-dependent dioxygenase**

OG253 is a protein possessing an Fe2+/2-oxoglutarate (2OG)-binding domain. Structural similarity analysis revealed homology to multiple Fe2+/2OG-binding proteins (e.g., PDB ID: 2RG4).

**OG271 Dipeptidyl peptidase**

OG271 is an uncharacterized gene annotated as COG3807. The encoded protein consists of a signal peptide and two SH3 domains, which are known to interact with proline-rich sequences (33). Structural comparison revealed similarity to putative dipeptidyl peptidase IV or VI enzymes derived from Bacteroides species (PDB: 4R0K, 3NPF, 3PVQ). Dipeptidyl peptidase IV is an N-terminal protein-processing enzyme that specifically cleaves dipeptides from the N-terminus when the second residue is a proline or alanine (34).

**OG228 AcrB-like transporter**

OG228 is an uncharacterized transporter gene containing two MmpL domains, which are known membrane transport domains (35). On the SAR11 genome, the DNA helicase gene dnaB is located in close proximity. Structural similarity analysis revealed a high degree of homology with transporters containing MmpL3 domains (e.g., PDB: 7N6B). OG228 also shows partial sequence and structural similarity to the multidrug efflux transporter AcrB. AcrB functions as an inner membrane pump by forming a complex with the adaptor protein AcrA and connects to outer membrane channels such as TolC or OprM to mediate multidrug efflux (36). However, neither acrA nor tolC homologs are found near OG228, and another copy of acrB is located adjacent to acrA in the SAR11 genome. These observations suggest that OG228 may transport substrates distinct from those of AcrB.

**OG346 Hydroxylase**

Structural similarity analysis revealed that OG346 shares structural homology with multiple hydroxylases (e.g., PDB: 7V4O, 4P7W), suggesting that it functions as a hydroxylase.

**OG397 Lytic transglycosylase**

OG397 is an uncharacterized gene annotated as COG4764 and encodes a protein with a predicted signal peptide. Structural similarity analysis revealed homology with several lytic transglycosylases (e.g., PDB: 5O1J, 5OHU, 5O29). Lytic transglycosylases are enzymes that degrade peptidoglycan by cleaving free glycan chains, thereby preventing their accumulation in the periplasmic space (37). These enzymes are also known to create localized gaps in the peptidoglycan layer to make insertion of structures such as the flagellar basal body (38). COG4764 sequences have been detected in marine environments, suggesting that it represents a gene specific to marine microorganisms.

**OG430 DUF2155-containing tyrosine phosphatase**

OG430 is a gene annotated as COG4765 and contains the domain of unknown function DUF2155. A previous study that clustered marine bacterial genes based on abundance correlations across sampling sites suggested that DUF2155 belongs to a cluster of genes involved in catalyzing the cleavage of nitrogen-containing moieties from polyamines and purines (39). On the SAR11 genome, the NADH-ubiquinone oxidoreductase gene NDUFA12 was found close to OG430. Structural similarity analysis revealed that OG430 shares significant structural homology with several protein tyrosine phosphatases (e.g., PDB: 4GE5, 6KRX), suggesting that OG430 may function as a tyrosine phosphatase.

**OG486 α/β hydrolase**

OG486 is an uncharacterized gene annotated as K07018 and COG2945. Structural similarity searches yielded hits to approximately 900 protein structures, including PDB 2I3D, with many of the top-ranked structures belonging to the α/β hydrolase superfamily. The α/β hydrolase superfamily is characterized by a conserved fold consisting of an eight-stranded β-sheet flanked by several α-helices, but despite this common fold, its members exhibit diverse evolutionary origins and functions (40).

**OG559 Putative ribonucleotide reductase**

The structure showed similarity to multiple Vitamin B12-dependent ribonucleotide reductases registered in the AlphaFold DB, suggesting that it may function as a ribonucleotide reductase.

**OG580 Cell cycle regulator**

OG580 is an uncharacterized gene containing a zinc ribbon motif. Structural similarity analysis revealed that the C-terminal zinc ribbon domain is homologous to that of ZitP (PDB: 2NB9) from *C. crescentus*. ZitP is known to be involved in pilus biogenesis and motility, and it has been shown to regulate cell polarity through interaction with PopZ (41–43).

**OG601 Glycosidase**

OG601 is an uncharacterized gene annotated as K03796 and COG2992, known as Bax. The gene is located near a tryptophanyl-tRNA synthetase gene in the SAR11 genomes. Structural similarity analysis revealed that OG601 shares structural homology with a peptidoglycan glycosidase associated with the type VI secretion system (PDB: 4KT3) (44).

**OG660 Lipoprotein**

OG660 is an uncharacterized gene encoding a protein with a predicted signal peptide. Structural similarity analysis revealed homology with a lipid-binding protein (PDB: 3BDR), suggesting that OG660 may function as a lipoprotein.

**OG628 Prolyl hydroxylase**

OG628 is an uncharacterized gene with structural similarity to proteins containing a 2-oxoglutarate (2OG)- and $Fe^{2+}$-binding domain. Structural similarity analysis revealed homology with the eukaryotic 2OG-dependent prolyl hydroxylase (PDB 4NHY) (45), and the prokaryotic elongation factor Tu prolyl hydroxylase (PDB 3IW3). These enzymes are believed to share a common evolutionary origin (46). OG628 is therefore predicted to be a prolyl hydroxylase of the same origin.

**OG635 DUF3576-containing lipoprotein**

OG635 is an uncharacterized gene encoding a protein with a predicted signal peptide and the domain of unknown function DUF3576. Structural similarity analysis revealed homology with the outer membrane lipoprotein BamC (PDB 2YH6, 6LYS) (47).

**OG678 Methyltransferase**

OG678 is an uncharacterized gene annotated as COG2940. Structural similarity analysis revealed homology to multiple eukaryotic histone methyltransferases (e.g., PDB 7W67, 2W5Z). Recent studies have shown that some proteobacteria possess histone-like proteins (48); however, it remains unclear whether SAR11 harbors histone-like proteins.

**OG709 Lipoprotein**

OG709 is a gene of unknown function that contains a predicted signal peptide. Structural analysis revealed similarity to lipoproteins, such as the lipoprotein from *Pseudomonas aeruginosa* PAO1 (PDB 4DM5), suggesting that OG709 encodes a lipoprotein.

**OG714 HepC, heparinase II/III-like**

OG714 is an uncharacterized gene annotated as COG5360 (Uncharacterized conserved protein, heparinase superfamily). Notably, 75.8% of COG5360 homologs have been detected in marine environments. Structural similarity to the heparan sulfate lyase from *Pedobacter heparinus* (PDB 4MMI) suggests that OG714 may function as a HepC-like enzyme (49).

**OG715 OmpF1-like porin**

OG715 is a gene of unknown function lacking annotation in COG databases. In SAR11 genomes, OG715 is syntenically conserved with neighboring genes OG635 (encoding a DUF3576-containing lipoprotein) and OG701 (COG3786, L,D-peptidoglycan transpeptidase). Predicted structural similarity to the *E. coli* porin OmpF1 (PDB 7FDY) suggests that OG715 functions as an OmpF1-like porin.

**OG721 Lon protease**

OG721 is an uncharacterized gene annotated as K07157 and COG2802. Structural similarity analysis revealed that it shares structural homology with ATP-dependent serine protease Lon protease (PDB: 7CR9, 3LJC) (50).

**OG735 RuvC endodeoxyribonuclease**

OG735 is an uncharacterized gene without a corresponding annotation in the COG database. Predicted structural similarity to multiple endodeoxyribonucleases, including the protein represented by PDB 6JRF, indicates that OG735 may function as an endodeoxyribonuclease.

**OG768 Helicase loader DciA**

OG768 is a gene annotated as COG5389 and contains the DUF721 domain. Structural similarity analysis revealed homology to DciA (PDB 7YKM, 8A3V), a helicase loader protein that facilitates the loading of the DNA helicase onto the replication origin during replication initiation (51,52).

**OG795 Dehydrogenase**

OG795 is annotated as K03937 and shows sequence similarity to NDUFS4, a redox enzyme component of mitochondrial respiratory complex I. Structural similarity analysis revealed a high degree of homology with eukaryotic respiratory complex I-associated proteins (e.g., PDB 7ZDP, 7VXU).

**OG796 Acetyltransferase**

OG796 is annotated as K01726 and COG0663 and is predicted to be a homolog of YrdA in *E. coli*. Structural similarity analysis suggested homology not only to *E. coli* YrdA but also to multiple acetyltransferases (e.g., PDB 4N27). YrdA belongs to the γ-class of carbonic anhydrases but is known to have minimal carbonic anhydrase activity (53). YrdA is predicted to form a homotrimer, and it is expected that OG796 similarly assembles into a trimeric structure.

**OG814 Na+/solute symporter**

OG814 is a membrane protein gene of unknown function, annotated as COG0679 (YfdV). Structural predictions revealed similarity to known $Na^+$/solute symporters, such as SbtA and a $Na^+$/bile acid symporter (PDB 7CYF, 4N7W), suggesting that OG814 may function as a $Na^+$/solute symporter.

**OG824 Bile acid sodium symporter homolog**

OG824 is an uncharacterized membrane protein gene annotated as COG0385 (YfeH). Its predicted structure shows similarity to the bile acid sodium symporter from *Neisseria meningitidis* (PDB 3ZUX), suggesting that OG824 may be a homolog of this transporter.

**OG828 SurA-like chaperone**

OG828 is an uncharacterized membrane protein. Structural similarity analysis revealed homology to SurA-like chaperones (e.g., PDB: 3RGC). SurA is a chaperone that facilitates the folding of outer membrane proteins (54). The genomic neighborhood of OG828 includes SecG, a subunit of the Sec translocase, and triosephosphate isomerase TpiA on the SAR11 genomes.

**OG853 VipF acetyltransferase**

OG853 is an uncharacterized gene lacking functional annotation in the COG database. Predicted structural similarity to several acetyltransferases (e.g., PDB 6WQB) suggests that OG853 may encode an acetyltransferase.

**OG859 LptE lipoprotein**

OG859 is an uncharacterized gene containing a predicted signal peptide. In SAR11 genomes, it is consistently located near OG645 (COG0495, leucyl-tRNA synthetase). Structural similarity to LptE from Pseudomonas aeruginosa (PDB 8H1R) suggests that OG859 encodes a lipoprotein.

**OG868 Cell division-related protein FtsB**

OG868 is an uncharacterized membrane protein gene. In April 2023, the structure of FtsB, a protein component of the bacterial divisome complex with structural similarity to OG868, was released in the PDB (PDB ID: 8HHG) (16). In *E coli*, FtsB is an essential gene known to form the FtsBLQ complex, which localizes FtsI and FtsW to the Z-ring (55).

**OG862 Metalloprotease TldE**

OG862 was annotated as K03592 and identified as a gene belonging to the metalloprotease TldD family. Structural similarity analysis revealed homology to TldE (e.g., PDB IDs 5NJC, 3QTD). Based on the structural similarity and previous studies (30,56), this gene is predicted not to encode TldD but rather TldE, a protein that forms a heterodimer with TldD.

**OG879 Chaperone**

OG879 is an uncharacterized gene annotated as COG5452. Structural similarity analysis revealed homology to CBP3, a ubiquinol–cytochrome c chaperone (PDB: 6RWT) (57).

**OG883 Polyketide synthase SnoaL-like**

OG883 is an uncharacterized gene lacking functional annotation in the COG database. In SAR11 genomes, it is consistently located near OG866, which encodes the membrane anchor protein AprM. Structural similarity searches revealed homology to calcium/calmodulin-dependent protein kinase type II association domains (e.g., PDB 3GWR), suggesting that OG883 may adopt a similar fold to SnoaL-like proteins (58).

**OG900 Succinate dehydrogenase protein subunit E (SdhE)**

Structural similarity analysis revealed that OG900 is homologous to *E. coli* ygfY (PDB 1X6J), a protein involved in FAD assembly (SdhE) (59).

**OG908 FtsQ-like SPOR domain-containing protein**

OG908 is an uncharacterized gene containing a predicted signal peptide region. Structural similarity searches revealed homology to proteins containing a SPOR domain (e.g., PDB: 4AVR, 6I05), suggesting that OG908 may be a homolog of FtsQ, a protein involved in bacterial cell division.

#### 4. Function predicted by KEGG Orthology annotation (19 OGs)

These OGs are presumed to be functionally known genes for which information in the COG database has not been updated.

**OG74 Zinc-binding hydrolytic enzyme GloB**

OG74 was annotated with K01069 and COG0491, and was identified as the GloB gene, which is a zinc-binding hydrolytic enzyme known to reduce oxidative stress by hydrolyzing S-lactoylglutathione, a degradation product of methylglyoxal in *E. coli* (60,61).

**OG154 AzoR**

OG154 was annotated as K01118 and COG2249, and predicted as an FMN-dependent NADH azoreductase (62).

**OG212 Cobalamin (vitamin B12) synthase CobS**

OG212 was annotated with K09882 and found to be a CobS gene in the aerobic cobalamin biosynthesis pathway (63). In its genomic neighborhood, genes encoding CobT, which forms a complex with CobS during cobalamin synthesis, and the molecular chaperone DnaJ were conserved. Structural similarity searches revealed that OG212 shares a similar fold with AAA+ domain-containing ATPases (e.g., PDB entries 6L1Q, 5C3C). Previously characterized CobS proteins are also known to function as AAA+ ATPases (64).

**OG225 Cell cycle regulator GcrA**

OG225 was annotated as K13583 and COG5352, corresponding to the cell cycle regulator GcrA, which is well characterized in the model alphaproteobacterium Caulobacter crescentus (65). GcrA regulates the cell cycle by forming a complex with RNA polymerase II to activate transcription (66). Although SAR11 is known to exhibit morphological changes depending on nutrient availability (67), cell cycle-dependent morphological transitions like those observed in *C. crescentus* have not been reported. This suggests that the set of genes regulated by GcrA in SAR11 may differ from those in *C. crescentus.*

**OG365 Anomeric sugar kinases ampK**

OG365 was annotated as K07102 and COG3178, corresponding to the anomeric sugar kinase AmpK, which functions in the peptidoglycan precursor salvage pathway (68). Structural similarity analysis revealed that OG365 shares significant structural features with over 200 kinases, including those represented by PDB entries 3CSV, 4FEX, and 2QG7.

**OG426 Ribosome small subunit protein RspQ**

OG426 was annotated K02961 and was rspQ, a ribosomal small subunit S17 (69). OG426 was encoded in the ribosomal protein operon.

**OG468 MidA family protein arginine methyltransferase**

OG468 was annotated as K18164 and COG1565 and identified as a SAM-dependent methyltransferase gene of the MidA family with previously unknown function. Structural similarity analysis revealed that it shares structural features with several methyltransferases, including MidA (PDB 5ZZW), supporting its classification as a methyltransferase. MidA is thought to be a methyltransferase required for the function of the mitochondrial respiratory chain complex (70). COG1565 is broadly conserved across Proteobacteria (71) and has been shown to methylate protein arginine residues via a mechanism similar to that of the mitochondrial methyltransferase NDUFAF7 (72).

**OG497 SecB**

OG497 was annotated as K03071 and was found to be the protein translocation-related gene secB (73).

**OG530 Chitin deacetylase**

OG530 was annotated as K21478 and identified as a chitin deacetylase gene belonging to the PgdA/NodB/CDA1 family. Structural similarity searches yielded over 600 hits, primarily to deacetylase enzymes, including those with PDB entries 6DQ3 and 4WCJ.

**OG592 Ribosome large subunit protein L23 RplW**

OG592 was annotated as K02892 and predicted as the gene encoding the large ribosomal subunit protein L23. In SAR11 genomes, it was also found to be encoded within the ribosomal protein operon.

**OG653 PagL**

COG653 was annotated as K12976 and predicted as PagL, an acyloxyacyl hydrolase involved in Lipid A biosynthesis (74).

**OG687 UDP-2,3-diacylglucosamine hydrolysis enzyme**

OG687 was annotated as K09949 and COG3494. In *C. crescentus*, it is located within an operon involved in membrane biosynthesis and has been shown to function in an alternative pathway for Lipid A synthesis (75). In SAR11 as well, it was found within a membrane biosynthesis-related operon.

**OG689 Ribosome large subunit protein L33 RpmG**

OG689 was annotated with K02913 and was found to be the ribosomal large subunit protein L33.

**OG717 Phosphoriblokinase**

OG717 is annotated as COG4240 and K15918, and predicted as a phosphoriblokinase.

**OG724 Quercetinase**

OG724 was annotated as K06911 and COG1741, and predicted as the Pirin gene, a quercetinase (76).

**OG783 Cell chape-determining protein MreD**

OG783 is annotated as K03571 and is predicted to correspond to mreD, a cell shape-determining gene (77). Approximately 93% of the sequences were detected in marine environments.

**OG840 DNA photolyase**

OG840 was annotated as K06876 and COG3046, and identified as a DNA photolyase gene (78,79). Structural predictions confirmed the presence of an iron-sulfur cluster-binding domain.

**OG856 Beta-carotene 15,15'-dioxygenase (blh)**

OG856 is annotated as K21817 and predicted to be the blh gene, which synthesizes retinal from beta-carotene. In SAR11 genomes, the neighboring OGs OG827 (COG1562, phytoene/squalene synthase ERG9) and OG860 (COG0665, glycine/D-amino acid oxidase DadA) are conserved.

### 5. Function Unknown (38 OGs)

No functional information could be obtained for these OGs.

**OG171 Uncharacterized membrane protein**

OG171 is an uncharacterized membrane gene annotated as COG3748. No structurally similar proteins were found in the PDB, and no conserved neighboring genes were identified.

**OG311 DUF1178-containing protein**

OG311 is a gene annotated as COG5319 and contains the domain of unknown function DUF1178. No structurally similar proteins or conserved neighboring genes were identified.

**OG327 DUF4167-containing protein**

OG327 is a gene containing the domain of unknown function DUF4167. The predicted structure consists of a hairpin-shaped α-helix flanked by disordered regions, and no structurally similar proteins were identified. An operon including the translation termination factor hemK (80) was conserved in the genomic vicinity.

**OG305 IalB-like membrane-pore protein**

OG305 is a functionally uncharacterized gene. Structural similarity analysis revealed that OG305 shares a monomeric structure similar to IalB, an outer membrane protein of unknown function required for infection of *Bartonella bacilliformis*, a causative agent of cat-scratch disease (81,82). IalB is registered in the PDB as a homo-dodecamer (PDB: 3DTD), suggesting that OG305 may also exist as a multimeric complex. To investigate this, we predicted the dodecameric structure of OG305 using AlphaFold2-Multimer, which produced a complex structure distinct from that of IalB in the PDB (see figure below). Based on the predicted structure and presence of a signal peptide, OG305 is likely to form a pore in the cell membrane as a homo-hexamer or dodecamer. Although the neighboring gene encoding argininosuccinate synthetase (ArgG) was conserved on the SAR11 genomes, its functional relationship to OG305 remains unclear.

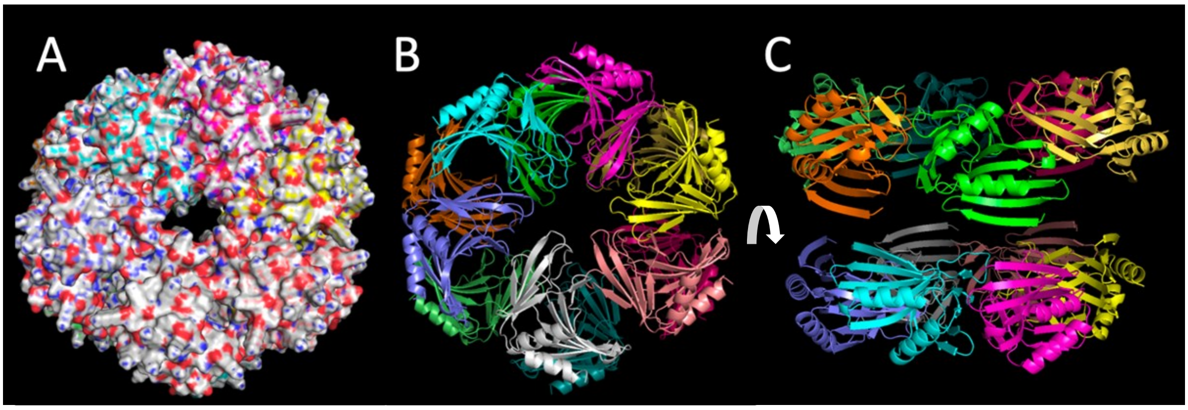


**Predicted structures of OG305 12-mer** (same as Fig.S8ABC)**.**

**OG251 Membrane protein of the DedA family, potentially associated with aerobic metabolism**

OG251 is annotated as COG0398, a putative membrane protein belonging to the DedA superfamily with unknown function. The DedA superfamily comprises three subfamilies (COG0586, COG0398, and COG1238). COG0398 lacks the conserved residues responsible for the UndP flippase activity found in the functionally characterized DedA subfamily COG0586. It is widely distributed among aerobic bacteria, and gene fusions with genes involved in aerobic metabolism—such as glutathione-thioredoxin reductase and lipoamide dehydrogenase—have been reported (17). COG0398 has been identified as an essential gene in model alphaproteobacteria such as *C. crescentus* and *Dinoroseobacter shibae* (83). In the SAR11 clade, the genomic region surrounding OG251 is conserved and includes the RNase P protein component RnpA.

**OG382 Function unknown periplasmic protein**

OG382 is an uncharacterized gene predicted to encode a protein with a signal peptide. In its genomic vicinity, genes encoding 3-dehydroquinate synthase and shikimate kinase I—components of the Escherichia coli dam superoperon—were found to be conserved (84). No structurally similar proteins have been identified in the PDB.

**OG535 Function unknown protein**

OG535 genomic neighborhood is conserved with OG525 (COG0717, dCTP deaminase) and OG602 (an uncharacterized protein containing a DUF2948 domain) in the SAR11 genomes, although any functional relationship between them remains unclear.

**OG543 Function unknown inner membrane protein FxsA**

OG543 is annotated as K07113 and COG3030, and has been identified as FxsA. FxsA is a non-essential inner membrane protein originally discovered in Escherichia coli for its role in suppressing F exclusion by bacteriophage T7 (85,86). However, its precise function remains unknown (87). In SAR11, the gene is located near secB, a gene involved in protein translocation (73). Based on the presence of a signal peptide and predicted transmembrane helices, FxsA in SAR11 is also predicted to be a membrane-associated protein. No structurally similar proteins have been registered in the PDB.

**OG590 Uncharacterized protein**

No structurally similar proteins to OG590 have been registered in the PDB, and its neighboring genes are not conserved. OG590 exhibits a four-helix bundle structure resembling that of the histidine kinase LytSN (88).

**OG602 DUF2948-containing protein**

G602 is an uncharacterized protein containing a DUF2948 domain. Structural similarity search revealed that its structure has been resolved for a sequence derived from *Neisseria gonorrhoeae* (PDB ID: 5V77); however, no functional information could be obtained. In its genomic neighborhood, dCTP deaminase (Dcd) and other genes of unknown function are conserved.

**OG669 DUF3108 or DUF6134-containing beta-barrel-like protein**

It contains the domains DUF3108 or DUF6134, and predicted structures indicate that it is a beta-barrel protein; however, its function remains unknown.

**OG690 Uncharacterized protein**

OG690 is a functionally uncharacterized gene possessing a signal peptide. The replication initiator gene DnaA is conserved in its genomic neighborhood.

**OG697 DUF3553-containing protein**

OG697 is an uncharacterized gene containing a DUF3553 domain, with no assigned COG or KO annotations. No conserved neighboring genes or structurally similar proteins were identified. The DUF3553 domain is predicted to adopt an SH3 β-barrel fold, and has been suggested to interact with RNA polymerase (89), although the function of this gene remains unknown.

**OG706 DUF2093-containing protein**

OG706 is annotated as COG3791, an uncharacterized conserved protein. It contains a DUF2093 domain, and its genomic neighborhood includes OG446 (COG1560; palmitoleoyl-ACP: Kdo2-lipid-IV acyltransferase LpxP), OG579 (COG0151; phosphoribosylamine–glycine ligase PurD), and

OG707 (COG1663; lipid A 4'-kinase LpxK), suggesting a potential role in lipid or purine biosynthesis, though its precise function remains unclear.

**OG730 DUF2794-containing protein**

OG730 is a functionally uncharacterized gene containing the DUF2794 domain. A heat shock protein HspQ gene is conserved in its genomic neighborhood.

**OG733 DUF2065-containing membrane protein**

OG733 is a functionally uncharacterized inner membrane protein that is annotated as K09937 and COG3242. It contains the DUF2065 domain, and its homolog is named YjeT in *E. coli*.

**OG754 DUF6151 super family CENP-V/GFA domain-containing protein**

OG754 is an uncharacterized gene annotated as COG3791 and belongs to the DUF6151 superfamily. It contains a CENP-V/GFA domain and shows structural similarity to an uncharacterized protein of *Rhodobacter sphaeroides* (PDB 3FAC).

**OG798 Function unknown protein PseE**

OG798 is annotated as K13256 and is predicted to be the phosphate starvation-induced gene PsiE. PsiE is related to a two-component system, but its exact function remains unknown (90,91).

**OG792 DUF2805-containing protein**

OG792 is a functionally uncharacterized gene containing the domain of unknown function DUF2805. No structurally similar proteins are registered in the PDB, and neighboring genes are not conserved.

**OG799 function unknown membrane protein**

OG799 is an uncharacterized gene predicted to encode a protein with transmembrane helices. In its genomic neighborhood, OG569 (COG2137; regulatory protein RecX) and OG907 (uncharacterized) are conserved, but no functional insights could be inferred from these associations.

**OG804 DUF2237-containing protein**

OG804 is annotated as K09966 and COG3651, containing a domain of unknown function DUF2237 (hereafter referred to as DUF2237). It is known that bacteria in the marine *Flavobacteriales* order tend to possess DUF2237 when they carry the gene for the light-driven proton pump proteorhodopsin. Knockout of DUF2237 in *Synechocystis* sp. PCC 6803-P strain results in reduced phototaxis (92). Approximately half of surface marine bacteria utilize light energy via proteorhodopsin, and light-utilization-related genes are crucial for understanding the ecology of marine bacteria (93). Therefore, we conducted a detailed functional analysis of DUF2237, a marine-specific gene related to light utilization.

**OG805 Uncharacterized protein**

OG805 is an uncharacterized gene annotated as COG4274 and contains a proline-binding domain known as GYD (94).

**OG823 DUF502-containing membrane protein**

OG823 was annotated as COG2928 and is a membrane protein gene containing the domain of unknown function DUF502. Approximately 74% of COG2928 sequences were detected from marine environments. DUF502 is a domain found in proteins involved in auxin transport in plants (95).

**OG835 Function unknown protein YqeY**

OG835 was annotated as K09117 and COG01610, corresponding to the yqeY gene. YqeY has been confirmed to be involved in the function of glutaminyl-tRNA synthetase (96), but it is also conserved in species lacking glutaminyl-tRNA synthetase, suggesting that it may have an unknown function (97).

**OG843 TPR-like protein**

OG843 is an uncharacterized gene predicted to encode a protein with a signal peptide. It shows structural similarity to PDB entry 1NA0, suggesting that it may contain a tetratricopeptide repeat (TPR) fold, which is commonly associated with protein–protein interaction.

**OG850 DUF6552-containing membrane protein**

OG850 is a membrane protein of unknown function containing the domain of unknown function DUF6552.

**OG863 DUF2721-containing membrane protein**

OG863 is an uncharacterized membrane protein gene containing a DUF2721 domain. In *Shewanella oneidensis*, point mutations in a membrane protein containing the DUF2721 domain have been reported to confer resistance to formate. However, docking simulations with formate indicate low binding affinity, and the function of this protein remains unknown (98).

**OG872 Uncharacterized membrane protein**

OG872 is a membrane protein gene of unknown function. It contains the low-oxygen-induced mitochondrial protein domain HIG1 (99), but no functional information was obtained.

**OG873 DUF4396-containing membrane protein**

OG873 is a membrane protein gene of unknown function containing the domain of unknown function DUF4396.

**OG875 DUF4864-containing protein**

OG875 is an uncharacterized gene predicted to encode a protein with a signal peptide. It contains a DUF4864 domain, the function of which remains unknown.

**OG880 BMFP Superfamily protein?**

OG880 is an uncharacterized protein group, with most of its sequences lacking COG and KO annotations. However, some sequences within this orthologous group are annotated as COG5493 (ubiquinone biosynthesis accessory factor UbiK), COG5493 (uncharacterized protein containing a PD-(D/E)XK nuclease domain), and COG0172 (seryl-tRNA synthetase), suggesting that OG880 may belong to the BMFP superfamily.

**OG886 Uncharacterized membrane protein**

OG886 is an uncharacterized gene predicted to encode a protein with transmembrane helices.

**OG894 DUF1467 family membrane protein**

OG894 is annotated as COG5454, an uncharacterized DUF1467 family protein found in Alphaproteobacteria.

**OG897 tetratricopeptide repeat-like domain-containing protein**

OG897 is an uncharacterized gene containing a tetratricopeptide repeat (TPR)-like domain. In its genomic neighborhood, OG725 (COG1520; outer membrane protein assembly factor BamB) is conserved, raising the possibility that OG897 may be functionally related to the BAM complex, although further evidence is required.

**OG904 uncharacterized protein (Potential esterase?)**

OG904 is an uncharacterized gene lacking functional annotation in the COG database. In SAR11 genomes, it is consistently located near OG936 (COG4327, uncharacterized membrane protein containing a DUF4212 domain). Although UniProt and the STRING database annotate OG904 as a potential esterase (e.g., Uniprot Q4FNV2), no structural similarity to known esterases was detected, and the basis for this annotation remains unclear.

**OG907 uncharacterized protein**

OG799 is conserved in the genomic neighborhood of OG907. However, both genes encode uncharacterized membrane proteins, and their functions could not be predicted.

**OG912 YGGT family membrane protein**

OG912 is a membrane protein of approximately 100 amino acids belonging to the YGGT family. In *E. coli*, YggT has been implicated in osmotic regulation, although the underlying mechanism remains unknown (100).

**OG918 Uncharacterized protein**

OG918 consists of very short sequences averaging 64 amino acids, and no functional information was obtained.
